# A Two-Component System FleS/FleR Regulates Multiple Virulence-Related Traits in *Pseudomonas aeruginosa*

**DOI:** 10.1101/2021.04.22.441042

**Authors:** Tian Zhou, Jia-hui Huang, Qi-shun Feng, Zhi-qing Liu, Qi-qi Lin, Zeling Xu, Lian-Hui Zhang

## Abstract

Microorganisms commonly use two-component systems (TCSs) to detect specific environmental changes and respond accordingly for their own benefit. However, the regulatory mechanisms and physiological roles of a majority of TCSs are still elusive. In this study, we focused on a previously predicted TCS FleS/FleR in *Pseudomonas aeruginosa* to systematically investigate its regulation and physiological roles. Loss of *fleS* or *fleR* or both genes led to decreased biofilm formation and attenuated motility in PAO1, which could be restored by heterologously complementation of FleR but not FleS, confirming that the sensor kinase FleS and the response regulator FleR constitute a TCS pair. To determine the regulatory spectrum of this TCS, we conducted transcriptome sequencing and comparison between the wild-type strain and the *fleR* deletion mutant. The result showed that the TCS regulates about 440 genes including most of them are involved in the virulence-related pathways, e.g. siderophore biosynthesis, pyocyanin biosynthesis, type III/VI secretion systems, c-di-GMP metabolism, flagellar assembly *etc*. In addition to its roles in controlling biofilm formation and motility we have already shown, FleR was demonstrated to regulate the production of virulence factors such as pyocyanin and elastase, mediate stress response to SDS, and autoregulate its own expression. Moreover, EMSA assays revealed that FleR regulates flagellum biosynthesis genes *flgBCDE*, *flgFGHIJKL*, *filC*, which are essential for the bacterial motility, by directly interacting with their promoters. Taken together, these results expanded our understanding on the biological roles of FleS/FleR and provided new insights on its regulatory mechanisms.

## INTRODUCTION

*Pseudomonas aeruginosa* is a ubiquitous Gram-negative bacterial pathogen which accounts for ~10% of hospital-acquired infections (Lyczak., 2003). This opportunistic pathogen frequently causes chronic lung infection, pulmonary inflammation, soft-tissue and other types of infections in immune compromised individuals (Deretic et al., 1995). *P. aeruginosa* can survive in diverse environments and outbreaks of drug-resistant strains are common among hospital wards and intensive care units (Costerton et al., 1995; Quinn., 2003). Human infections caused by *P. aeruginosa* is greatly attributed to its capabilities of producing various virulence factors, such as pyocyanin, elastase, rhamnolipids, exotoxins, lipopolysaccharides *etc*. (Dong et al., 2008; Wang et al.,2013). Moreover, swimming in liquid environments and swarming on semisolid surfaces are two major types of motility in *P. aeruginosa*, which enable the pathogen to expand the colonization niches and lead to systemic infections (Drake et al., 1988; Wang et al., 2014; Lai et al., 2009; Yeung et al., 2012). If not eradicated by the human immune systems, continuous infection of *P. aeruginosa* can result in its adaptation to human environment with biofilm formation, which increases its persistence and finally establishes long-term chronic infections (Costerton et al., 1995; Sousa et al, 2014).

The diverse virulence traits of *P. aeruginosa* are regulated by various regulatory systems such as two-component systems (TCSs), quorum sensing (QS) systems and host-pathogen cell-cell communication systems (Lee et al., 2015; Ahator et al., 2019). Bacterial TCS is one of the most common signal transduction systems with which bacteria perceive, respond and adapt to changes in the surrounding environment (Dong et al., 2008). A TCS typically consists of an inner transmembrane histidine sensor kinase and a response regulator with a signal receiver domain and a DNA binding domain. The sensor kinase detects environmental stimuli and autophosphorylates the conserved histidine residue of the kinase domain, which subsequently phosphorylates an invariant aspartate residue at the receiver domain of the cytoplasmic response regulator. The activated response regulator then regulates the expression of downstream genes via protein-DNA interaction (He et al., 2006 and 2009). Bioinformatics analysis identified approximately 64 sensor kinases and 73 response regulators in *P. aeruginosa* (Rodrigue et al., 2000; Galperin et al., 2006), indicating the pathogen had evolved sophisticated mechanisms to adapt to the changing environments. Among them, a few TCSs are known for regulation of various virulence traits in *P. aeruginosa*. For example, GacS/GacA is involved in regulating quorum sensing via small RNAs (Kay et al., 2006; Brencic et al., 2009), BqsS/BqsR influences rhamnolipids production and biofilm formation (Dong et al., 2008), and FimS/AlgR regulates alginate biosynthesis, motility and cytotoxicity (Deretic et al., 1989; Whitchurch et al.,1996). Despite these progresses, the biological roles and regulatory mechanisms of many other TCSs in *P. aeruginosa* have not yet been fully elucidated.

The genes *PA1098* and *PA1099* are predicted to encode a TCS, designated as FleS/FleR. Both FleS and FleR are found essential for swarming motility in the *P. aeruginosa* strain PA14 (Kollaran et al., 2019). Supporting its role in bacterial motility, mutation of *fleR* was shown to abrogate the biogenesis of flagellum in the *P. aeruginosa* PAK strain (Ritchings et al., 1995). In addition, previous studies showed that expression of FleS/FleR is regulated by FleQ and another TCS PilS/PilR (Jyot et al., 2002; Kilmury et al., 2018). Interestingly, the *fleS* and *fleR* deletion mutants displayed attenuated cytotoxicity against cultured human bronchial epithelial cells (Gellatly et al., 2018), suggesting that this TCS potentially contributes to the regulation of other virulence-related traits in *P. aeruginosa* in addition to motility. In this study, we first showed the role of the histidine kinase FleS and the response regulator FleR in biofilm formation and motility and then verified they constitute a TCS pair. Transcriptome and phenotypic analyses showed FleS/FleR regulates multiple phenotypes such as production of pyocyanin and elastase and mediates SDS response in addition to biofilm formation and motility. Finally, we presented evidence that FleR directly interacts with target gene promoters to autoregulate its own expression and control flagellum biosynthesis. This study presented a comprehensive investigation on the regulation and biological functions of the TCS FleS/FleR and provided insights on TCS-regulated virulence-related traits in bacterial pathogens.

## MATERIALS AND METHODS

### Bacterial strains and growth conditions

*P. aeruginosa* strains and other bacteria used in this study are listed in the supplementary information Table S1. Unless otherwise indicated, *P. aeruginosa* wild type and its derivatives, and *Escherichia coli* strains were routinely grown at 37°C in either Luria-Bertani (LB) broth (tryptone 10g/L, yeast extract 5g/L, NaCl 10g/L) or corresponding agar medium. Antibiotics were added when necessary at the following concentrations: gentamicin, 50 μg/ml for *P. aeruginosa* and *E. coli*; kanamycin, 50 μg/ml for *E. coli*; ampicillin, 100 μg/ml for *E. coli*. Bacterial cell density was determined by measuring optical density (OD) at the wavelength of 600 nm.

### Construction of mutants

The pK18mobsacB plasmid was used to construct in-frame deletion mutants of *P. aeruginosa* as previously described (Feng et al., 2020). The plasmids and primers used in this study were listed in Table S1 and Table S2, respectively. For instance, to generate the *fleS* gene deletion mutant, 500-bp upstream and 500-bp downstream homologous arms of *fleS* were amplified by PCR using a specific primer pair (Table S2) with Pfu DNA polymerase (Vazyme, China). After digestion with BamHI and HindIII, the PCR products were cloned into the suicide vector pK18mobsacB, generating pK18-*fleS* for *fleS* deletion. The resultant construct pK18-*fleS* was introduced into the *P. aeruginosa* strain PAO1 with the helper plasmid pRK2013 by triparental mating. Recovered colonies were selected and streaked on LB agar plates containing 10% sucrose. Desired colonies were selected by its susceptibility to gentamicin and tolerance to sucrose and further confirmed by PCR and DNA sequencing.

### Complementation analysis

For *in trans* complementation of mutants, the coding region of a gene was first amplified together with its native promoter from the PAO1 genome by PCR. The PCR product was cloned downstream of the *lac* promoter in the shuttle vector pBBR1-MCS5 after digestion by HindIII and BamHI. The resultant construct was verified by sequencing and then introduced into the corresponding PAO1 mutants by triparental mating. The complemented strains were confirmed by PCR analysis.

### Biofilm formation assay and quantification

Biofilm formation assay was performed according to the method previously described with minor modifications (An et al., 2010). Briefly, overnight bacterial cultures were diluted to an optical density at 600 nm (OD_600_) of 0.002 with fresh LB broth. The diluted cultures (150 μl) was transferred to 96-well polypropylene microliter plates and incubated at 37°C for the indicated periods of time. Bacterial cell density (OD_600_) was measured by a microplate reader (BioTek, USA). Bacterial cultures were carefully removed and the plates were washed three times with water. The biofilm cells bound to the walls of the plate were stained with 0.1% crystal violet (175 μl) for 15 min at room temperature, and then rinsed three times with water. The plates were air dried at room temperature. For quantification, biofilms were suspended in 200 μl of 95% ethanol and its absorbance at 570 nm was measured with a microplate reader. The concentration values were normalized to the cell density of each sample (OD_570_/OD_600_). All experiments were performed three times with six replicates.

### Motility assays

The motility assays were performed as described previously by Rashid et al. (Rashid et al. 2000) with minor modifications. For swimming motility, tryptone medium (10 g/L Tryptone, 5 g/L Yeast extract, 0.25 g/L Agar) was used. Swimming plates were dried at room temperature for about 10 min in a biosafety cabinet and then inoculated with 1 μl bacterial cells from an overnight culture grown in LB broth at 37°C. The plates were then wrapped with Saran Wrap to prevent dehydration and incubated at 37°C for 14 h before measuring of motility. The medium used for swarming assay consists of 0.5% (wt/vol) agar, 8 g/L nutrient broth and 5 g/L glucose. Swarming plates were typically allowed to dry at room temperature for forty minutes before being used. Swarming plate was inoculated with 1 μl overnight bacterial culture and inoculated at 37°C for 16 h before measurement. All experiments were performed three times with triplicates.

### Pyocyanin quantitation assay

Pyocyanin concentration was determined as described by Welsh et al. (Welsh et al., 2015). Briefly, the bacterial cultures were centrifuged at 14,000 rpm after grown for 16 h in 3 ml of LB medium. The supernatants were collected and filtered to remove residue cells. The absorbance of the supernatant was measured at 695 nm. The concentration values were normalized to the cell density of each sample (OD_695_/OD_600_). The experiments were performed three times with triplicates.

### Elastase assay

Elastase activity was assayed by elastin-Congo red (Sigma) assay with minor modifications (Ohman et al., 1980). Briefly, *P. aeruginosa* and its derivatives were grown in 3 ml LB medium at 37 °C for 16 h with shaking at 200 rpm. 500 μl aliquot of bacterial supernatants was added to an equal volume of 5 mg/ml elastin-Congo red in ECR buffer and incubated for 3 h at 37 °C. The amount of Congo red dye released from the digested elastin was determined using a spectrophotometer at A520, which is proportional to the activity of elastase in the supernatant. The activity values were normalized to the cell density of each sample (OD_520_/OD_600_). All experiments were performed three times with triplicates.

### RNA extraction, RT-PCR and quantitative real-time PCR

*P. aeruginosa* and its derivatives were grown in LB medium until the OD_600_ reached 1.5. Total RNA was isolated using the RNeasy mini kit (Qiagen, Germany) according to the manufacturer’s instructions. The cDNA samples were synthesized from the isolated total RNA using SuperScript II reverse transcriptase (Invitrogen, USA) and random primers (Invitrogen, USA). RT-PCR was performed on ProFlex™ PCR (Thermo Fisher Scientific, USA) using DNA polymerase (QingkeBiotech, China). The same corresponding batch of cDNA was used for quantitative real-time PCR (qRT-PCR) analysis. qRT-PCR was performed using the QuantiTect SYBR Green PCR kit (Qiagen, Germany) on the ABI QuantStudioTM^6^ Flex system (Roche, Switzerland) according to the manufacturer’s instructions. The primers used in this experiment were listed in Table S2. The experiment was repeated three times with triplicates.

### RNA-seq analysis

The enriched mRNA was fragmented as 200-700 nt and reverse transcribed into cDNA with random primers. Second-strand cDNA was synthesized by DNA polymerase I, RNase H, dNTP, and buffer. Then the cDNA fragments were purified with QiaQuick PCR extraction kit with end repaired and poly (A) added and ligated to Illumina sequencing adapters. The ligation products were size selected by agarose gel electrophoresis, followed by PCR amplification, and sequencing by Illumina HiSeq TM 2500 (Gene Denovo Biotechnology Co., China). Differentially expressed genes with ≥ 1.2-Log2fold changes were identified at a false discovery rate (FDR) ≤ 0.05, and analyzed using the major public pathway-related database KEGG (Kanehisa et al., 2008). The calculating formula for *p* value is

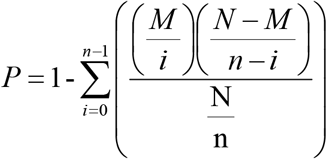

where N is the number of all genes that with annotation in database, n is the number of differentially expressed genes in N, M is the number of all genes annotated to specific pathways, and m is the number of differentially expressed genes in M. The calculated *p* value was gone through FDR correction, taking FDR ≤ 0.05 as a threshold. Q value is the *p* value underwent multiple hypothesis test corrections. The value ranges from 0 to 1 with more significant when it is closer to 0.

### SDS-induced macroscopic aggregation

The experiment was performed as described previously (Chen et al., 2020). Briefly, overnight cultures of bacterial strains were diluted (1:100) into 1.5 ml M9 medium (6.78 g/L Na_2_HPO_4_, 3 g/L KH_2_PO_4_, 0.5 g/L NaCl, 1.0 g/L NH_4_Cl) containing 3.5 mM SDS and cultured in 12-well plates with shaking at 120 rpm for 18 h at 30°C. Cell aggregation was quantified by measuring the size of bacterial aggregation. All experiments were performed three times with triplicates.

### Protein expression and purification

The open reading frame of *fleR* was amplified with the primers listed in Table S2, and subcloned into the expression vector pGEX-6p-1. The resulting construct was transformed into *E. coli* strain BL21 for FleR expression. The glutathione-Sepharose 4B beads (Smart life, China) were used for the purification of the GST-FleR fusion following the methods described previously (An et al., 2010). GST-tag cleavage was conducted with PreScission Protease (SMART Lifesciences, China; 2 units/ μl of bound proteins) at 4°C for 16 h. The obtained FleR protein was collected and analyzed by SDS-PAGE.

### Electrophoretic gel mobility shift assay

The DNA probes used for electrophoretic gel mobility shift assay (EMSA) were prepared by PCR amplification using the primer pairs listed in Table S2. The purified PCR products were 3’-end-labelled with biotin following the manufacturer’s instruction (Thermo Fisher Scientific, USA). The DNA-protein binding reactions were performed according to the manufacturer’s instructions (Thermo Fisher Scientific, USA). The 4% polyacryl gel was used to separate the DNA-protein complexes. After UV cross-linking, the biotin-labeled probes were detected in the membrane using a biotin luminescent detection kit (Thermo Fisher Scientific, USA). The EMSA experiment was performed three times.

### Statistical analysis

Experimental data were analyzed by one-way analysis of variance (ANOVA) and means were compared by Bonferron’s multiple comparison test using Graphpad Prism software (version 8). Experiments were arranged as completely randomized design and differences at *p* <0.05 were considered as statistically significant.

## RESULTS

### FleS and FleR are involved in the regulation of biofilm formation and motility

To understand the role of FleS/FleR in regulating bacterial virulence, we generated the *fleR* and *fleS* in-frame deletion mutants of *P. aeruginosa* strain PAO1. We first evaluated whether deletion of *fleS* and *fleR* would affect bacterial growth. We measured growth curves of the wild-type PAO1, Δ*fleS*, Δ*fleR* and their corresponding complemented strains and the result showed that there is no significant difference among these strains (Fig. S1). Next, a time-course analysis of biofilm formation was performed over a period of 30 h (Fig. 1A). It was shown that the biofilm biomass of the wild-type PAO1 was increased and reached a maximum amount at the time point of 10 h followed by a progressive decrease (biofilm decay). Compared to the wild type, deletion of *fleR* showed decreased biofilm biomass at all time points, whereas deletion of *fleS* produced same amounts of biofilm as the wild type at the first 6 h but its biofilm decay was observed about 4 h earlier. We then examined the impact of *fleS* and *fleR* on cell motility. The result showed that the swarming motility was reduced substantially in both Δ*fleS* and Δ*fleR* strains, however, deletion of *fleS* only moderately reduced swimming motility which was also substantially reduced by the deletion of *fleR* (Fig. 1B, 1C). *In trans* expression of wild-type *fleS* and *fleR* in the corresponding mutants rescued biofilm formation, swarming and swimming motility to wild-type levels (Fig. 1A-1C). These results indicated that FleS and FleR are involved in modulating biofilm formation and motility, and FleR plays more critical role than FleS. These findings were similar but not identical to the results of a previous study which showed that deletion of either *fleS* or *fleR* in PAO1 led to significant reductions in both swimming and swarming motility (Gellatly et al., 2019). We speculated that such differences may be due to the genetic dissimilarity occurred in different PAO1 sublines (Klockgether et al., 2010).

**Fig. 1.**
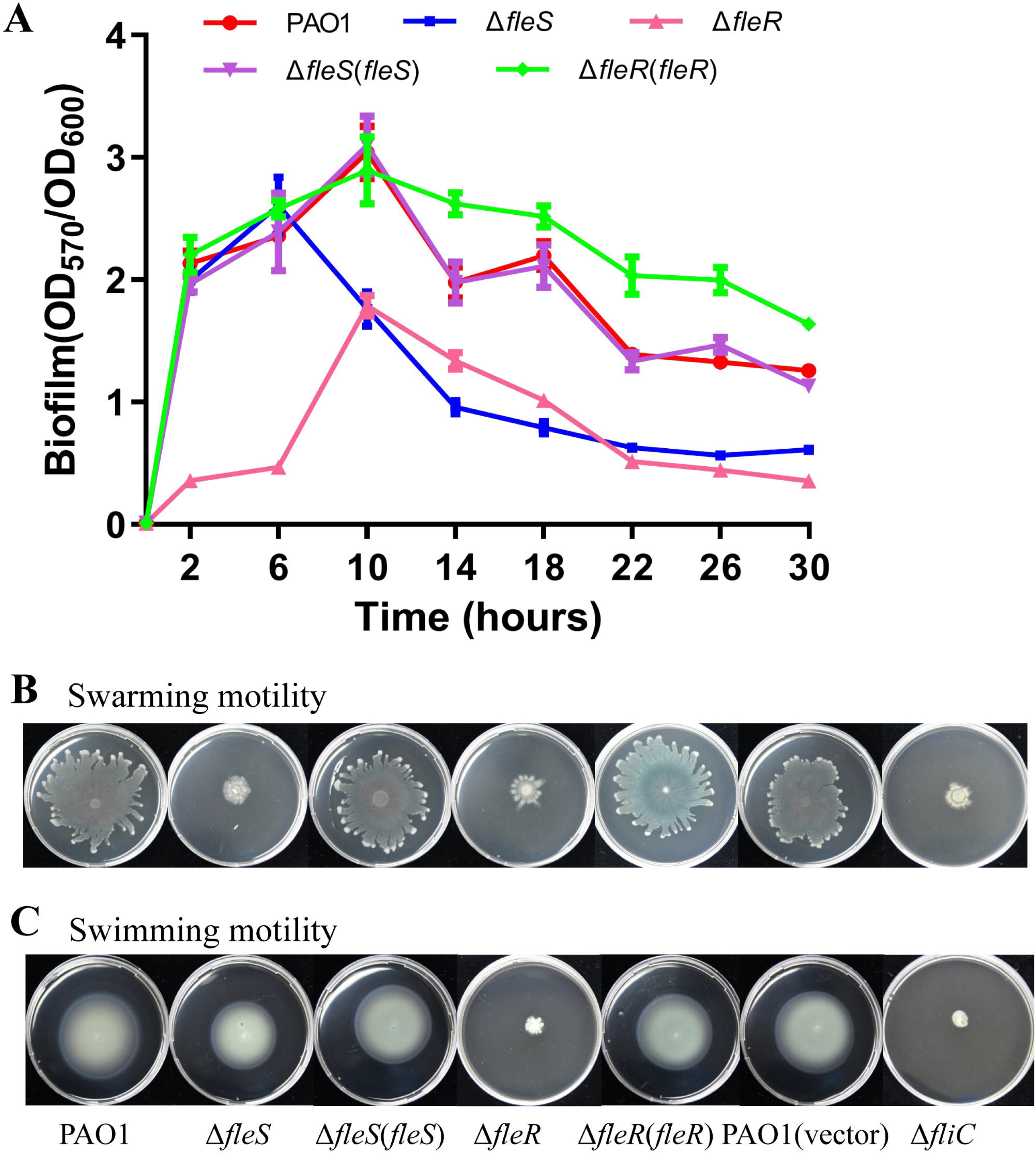
FleS and FleR regulate biofilm formation and motility in PAO1. (**A**) Time-course analysis of biofilm formation in PAO1 and its *fleS* or *fleR* mutants as well as their complemented strains. The data is the mean of six replicates with standard deviations. (**B**) Swarming and (**C**) swimming motility of PAO1 and its *fleS* or *fleR* mutants as well as their complemented strains. The vector pBBR1-MCS5 in wild-type PAO1 serves as the empty control for complemented vectors and the *fliC* mutant serve as the negative control for the motility assay.

### FleS and FleR constitute a two-component regulatory system

In PAO1, *fleS* and *fleR* are separated only by 4 base pairs and transcribed with the same orientation (Fig. 2A), suggesting that they are transcriptionally coupled and functionally related. Domain structure prediction using SMART program (http://smart.embl-heidelberg.de/) showed that the *fleS* gene encodes a protein with 402 amino acids, containing a PAS domain, a histidine kinase domain, and a histidine kinase-like ATPase domain. However, unlike typical TCS sensor kinases, the predicted FleS lacks a transmembrane domain (Fig. 2A). Different with the ubiquitous response regulators which contain REC-HTH domains only, FleR is relatively large in size with 473 amino acids containing REC-AAA-HTH domains (Fig. 2A). Reverse transcription polymerase chain reaction (RT-PCR) analysis confirmed that these two genes belong to the same operon (Fig. 2B).

**Fig. 2.**
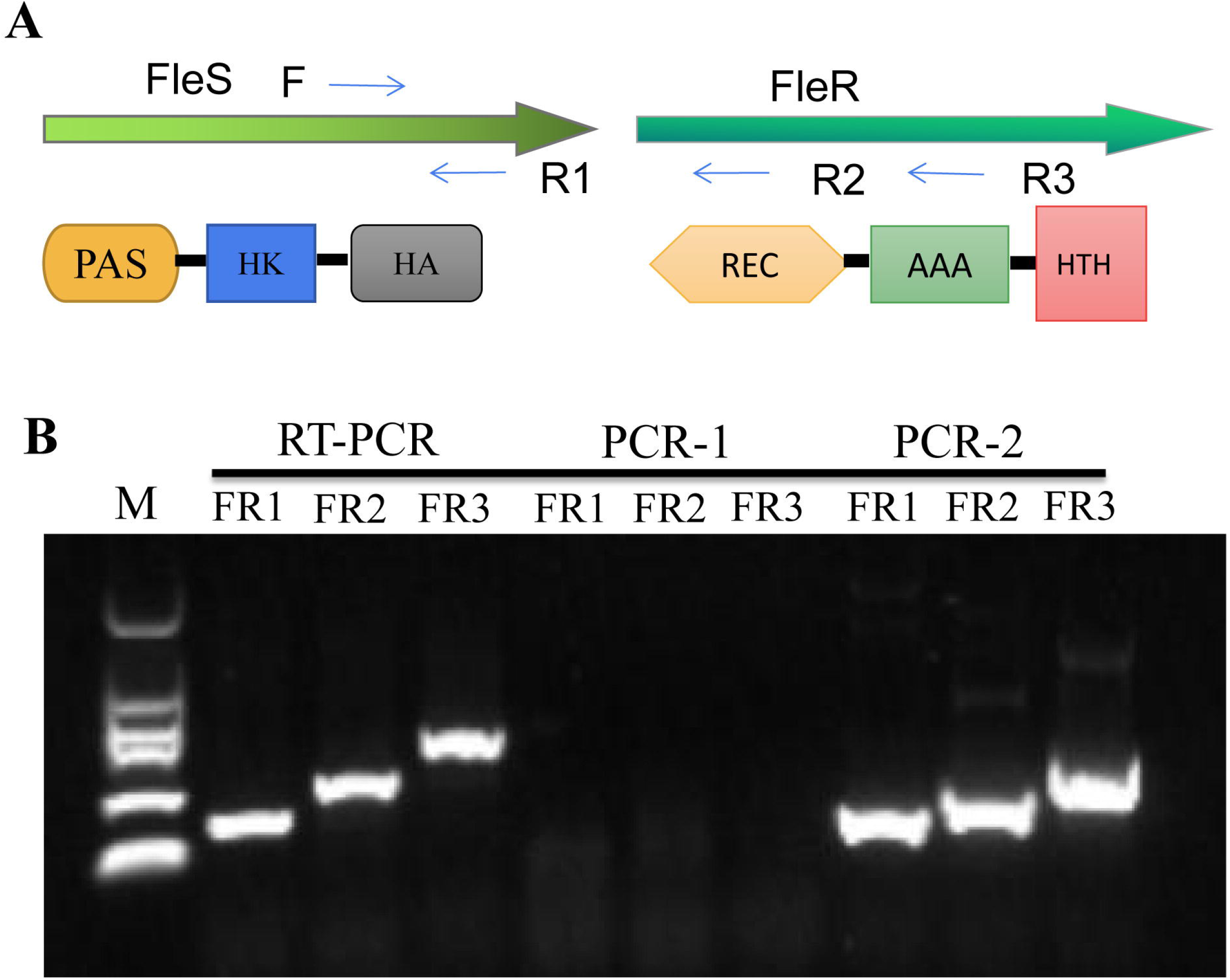
Analysis of the *fleSR* operon and the domain structures of their products. (**A**) Predicted genetic organization of the sensor kinase gene *fleS* and the response regulator gene *fleR* with their transcriptional orientation are indicated by open arrow. The relative locations of primers F1, R1, R2 and R3 used for RT-PCR analysis are indicated by solid arrows. The panels below are the domain structures of FleS and FleR, respectively, predicted by the SMART program. PAS: PAS domain; HK: histidine kinase A domain; HA: histidine kinase-like ATPase domain; REC: CheY homologous receiver domain; AAA: AAA domain; HTH: helix-turn-helix domain. (**B**) RT-PCR analysis based on the cDNA sample showed that *fleS* and *fleR* are co-transcribed. The RNA sample used for cDNA synthesis is used for PCR-1 analysis to preclude the possibility of genomic DNA contamination in the purified total RNA samples. Genomic DNA of PAO1 is used as the template for PCR-2 as a positive control.

To further confirm *fleS* and *fleR* are TCS pairs, we generated heterologously complemented strains Δ*fleR*(*fleS*) and Δ*fleS*(*fleR*) and examined its biofilm formation at 10 h and swimming motility. The results exhibited that *in trans* expression of *fleR* in Δ*fleS* but not *fleS* in Δ*fleR* restored its biofilm formation (Fig. 3A) and swimming motility (Fig. 3B). Moreover, we tested the *in trans* expression of *fleS* and *fleR* in the double deletion mutant Δ*fleS*Δ*fleR*, respectively, and the result showed that expression of *fleS* in the mutant Δ*fleS*Δ*fleR* failed to restore its capacity of biofilm formation and swimming motility while expression of *fleR* fully restored its biofilm formation and swimming motility to wild-type levels (Fig. 3A, 3B). Combined, these findings verified that FleS and FleR are TCS pairs and FleR is the cognate response regulator of the sensor kinase FleS.

**Fig. 3.**
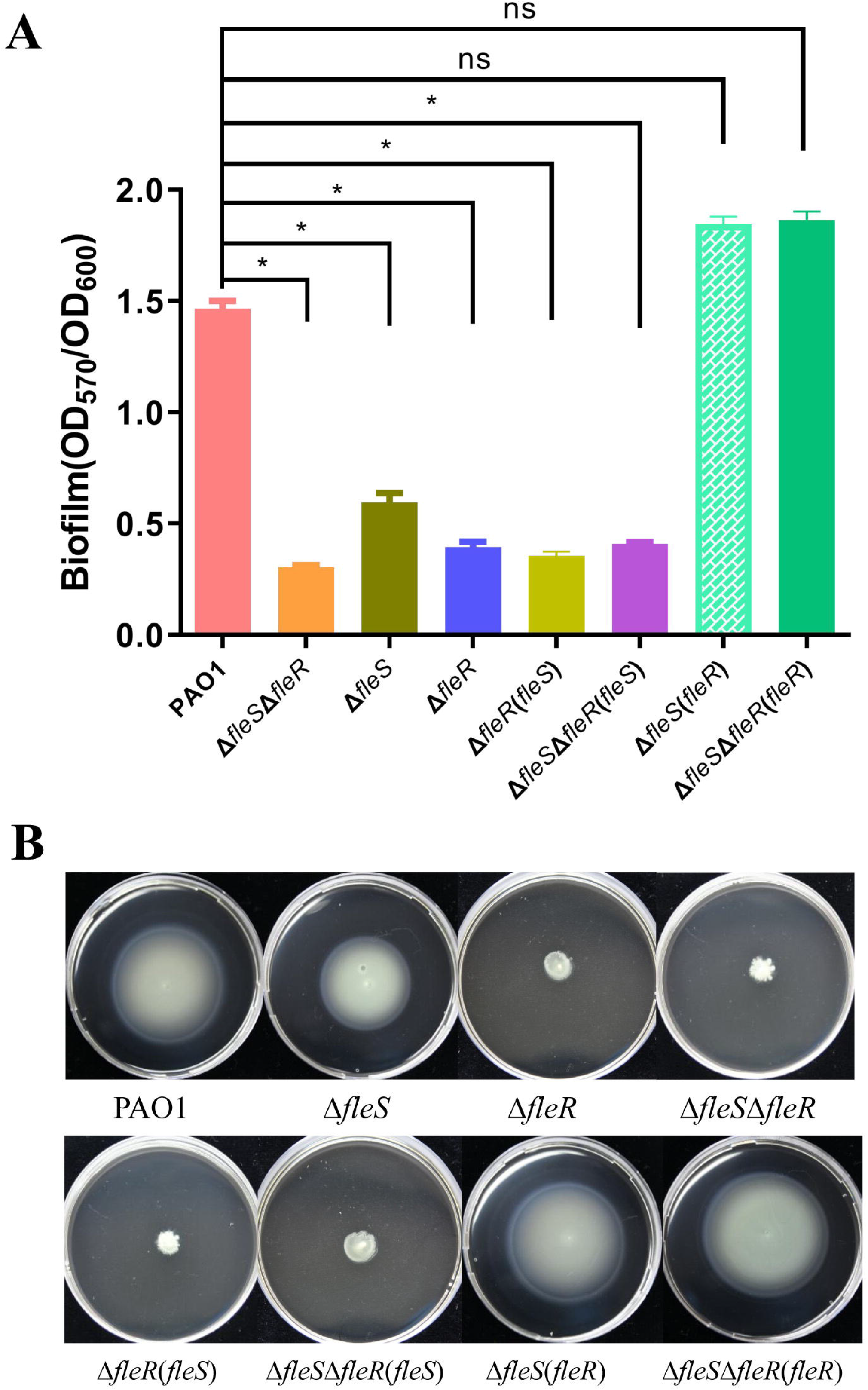
FleS and FleR constituted a two-component system. (**A**) Quantitative determination of biofilm formation of PAO1 and its derivatives after 10-h inoculation. The data is the mean of six replicates with standard deviations. *: *p*< 0.05, ns: not significant, tested by Student’s *t*-test. (**B**) Swimming motility of PAO1 and its derivatives.

### FleR controls the transcription of a wide range of genes belonging to diverse pathways in PAO1

In attempt to comprehensively understand the regulon of the TCS FleS/FleR, we analyzed and compared the transcriptomes of the wild-type PAO1 strain and its isogenic mutant Δ*fleR*. Considering that the obvious difference in biofilm formation of these two strains was observed between the time points 2 and 6 h (Fig. 1A), we collected the bacterial cells after 4-h growth for RNA-seq analysis. A total of 440 genes were identified with more than 1.2-log_2_fold changes in expression. Specifically, 121 genes were downregulated and 319 were upregulated by the deletion of *fleR* (Table S3). Overall, KEGG analysis revealed that the differentially expressed genes could be clustered into 20 functional groups (Fig. 4A). As summarized in Table 1, most identified genes with significant expression changes are virulence-related genes, e.g. genes involved in siderophore biosynthesis, pyocyanin biosynthesis, type III/VI secretion systems, c-di-GMP metabolism, flagellar assembly *etc*. Expression of these genes were subjected to verification by qRT-PCR (Fig. S2, S3). Given that flagellum and c-di-GMP are essential for motility and biofilm formation, respectively (Kearns et al., 2010; Merritt et al., 2007; Jones et al., 2014), the identification of flagellum biosynthesis genes and c-di-GMP metabolism genes in the RNA-seq result explains the connections of FleR with biofilm formation and motility.

**Fig. 4.**
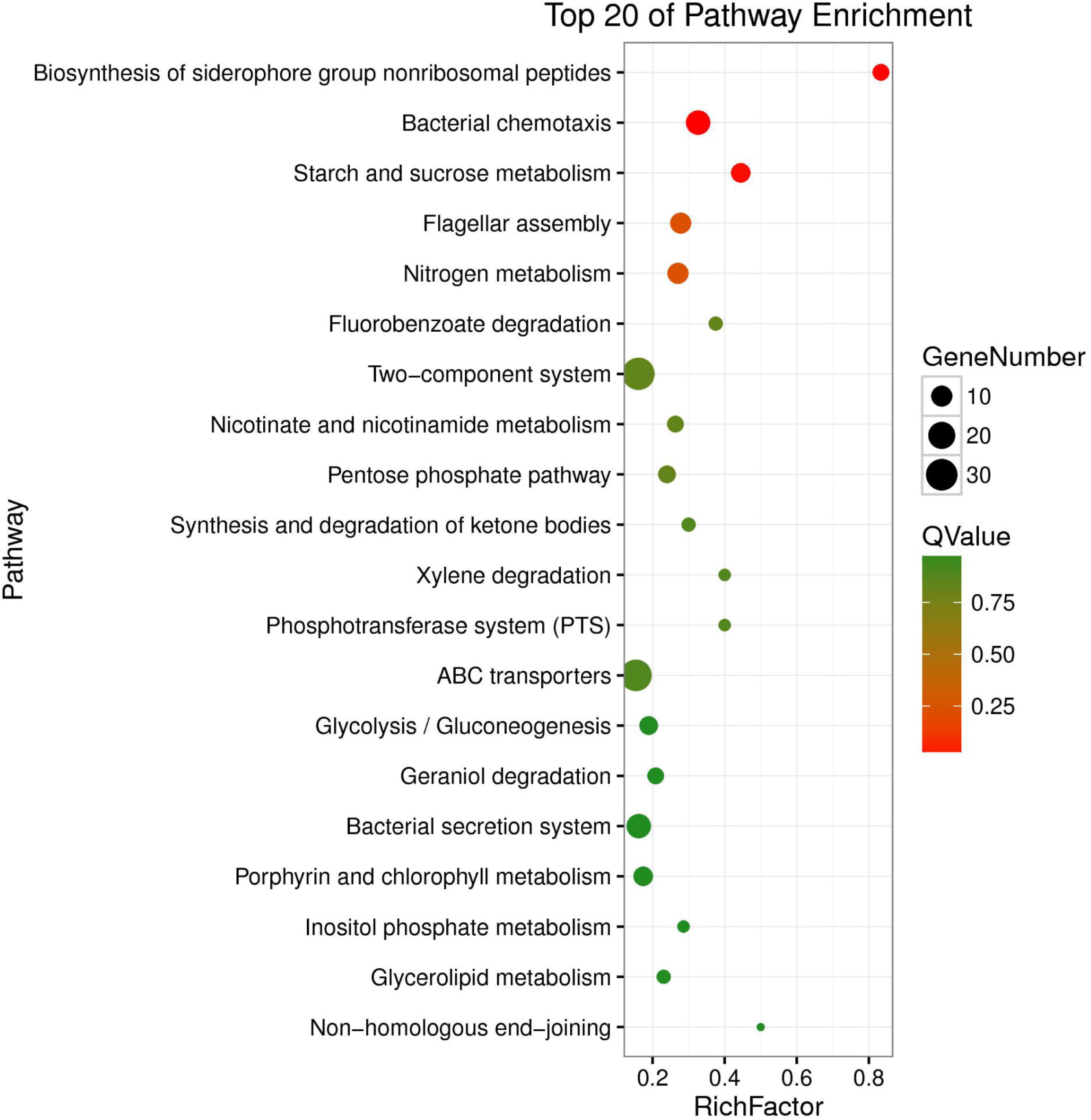
KEGG-enrichment of differentially expressed genes identified in RNA-seq analysis. The Y axis represents the names of the pathways. The X axis represents the rich factor. The size of the dot represents the number of differentially expressed genes in the pathways, and the color of the dot represents different Q values. The higher the value of rich factor represents the greater the enrichment degree. The smaller the Q value represents the more significant the enrichment. Rich factor index is used to measure the enrichment degree of pathway. Rich factor refers to the ratio of the number of genes annotated to the pathway in differentially expressed genes to the total number of genes in the pathway in all annotated genes.

**Table 1.**
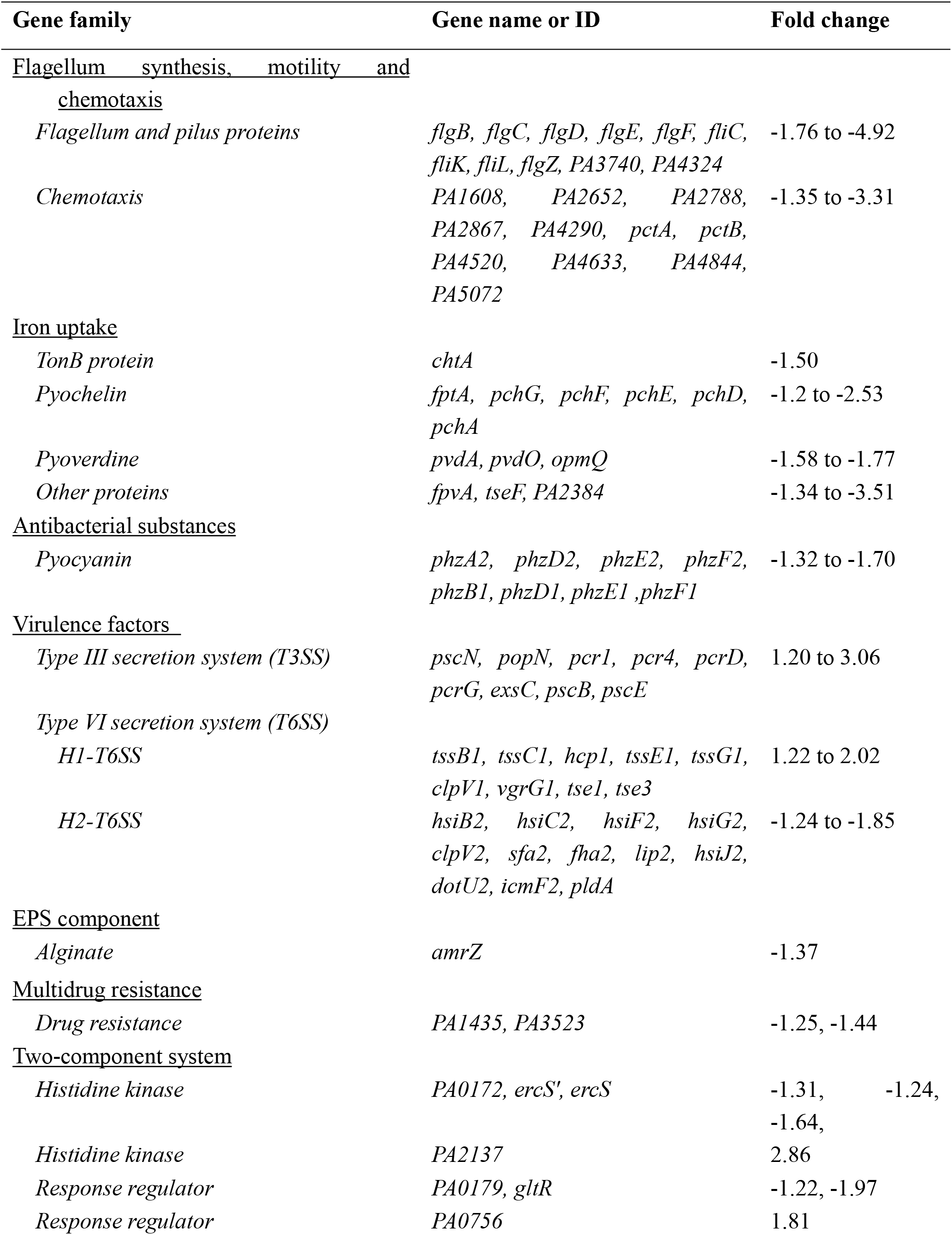
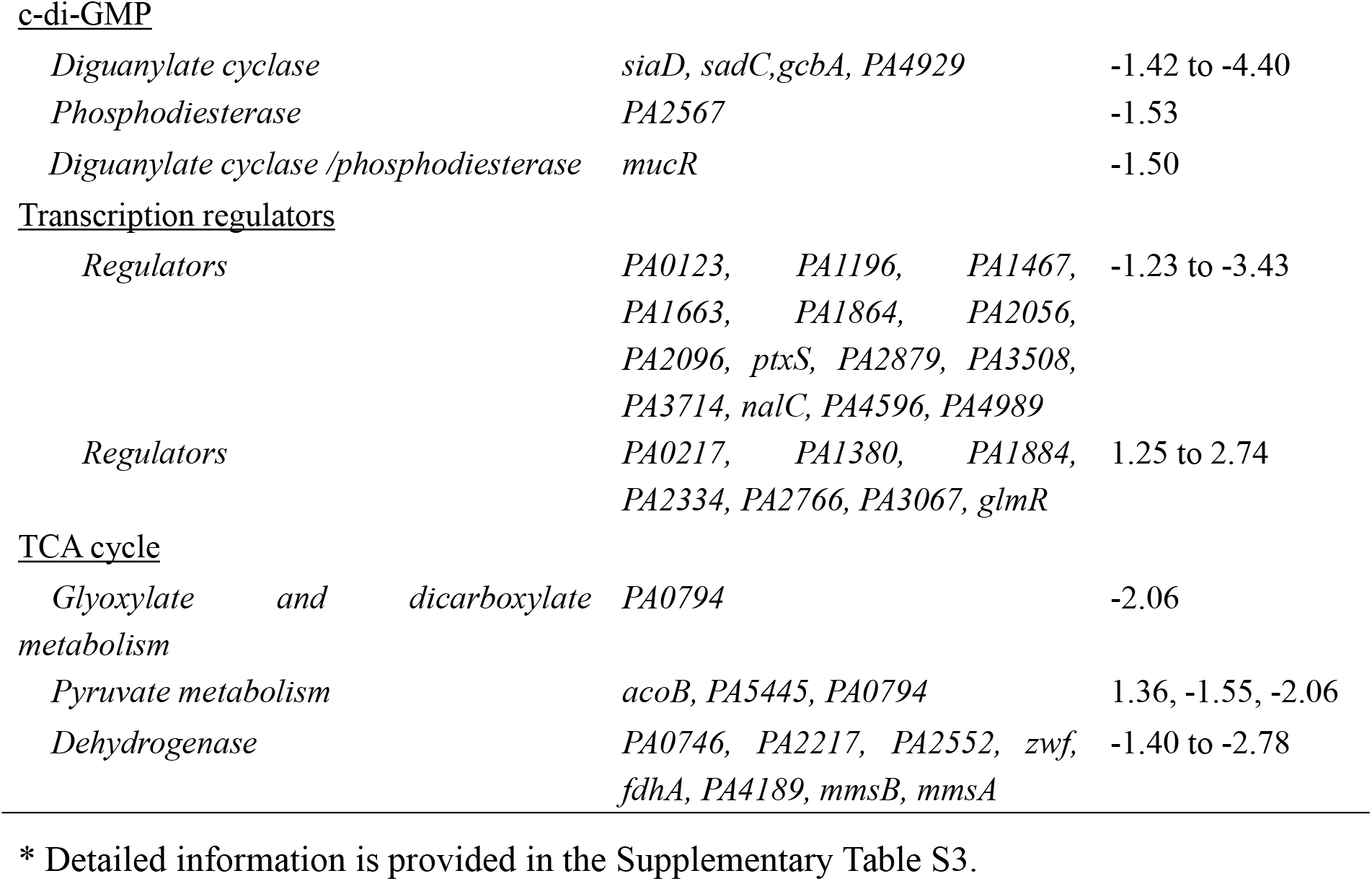
Selected gene families with more than 1.2-Log_2_fold changes owing to the deletion of *fleR* in PAO1 (Δ*fleR*/PAO1 WT)*.

| Gene family | Gene name or ID | Fold change |
| --- | --- | --- |
| <u>Flagellum synthesis, motility and chemotaxis</u> |  |  |
| *Flagellum and pilus proteins* | *flgB, flgC, flgD, flgE, flgF, fliC, fliK, fliL, flgZ, PA3740, PA4324* | -1.76 to -4.92 |
| *Chemotaxis* | *PA1608, PA2652, PA2788, PA2867, PA4290, pctA, pctB, PA4520, PA4633, PA4844, PA5072* | -1.35 to -3.31 |
| <u>Iron uptake</u> |  |  |
| *TonB protein* | *chtA* | -1.50 |
| *Pyochelin* | *fptA, pchG, pchF, pchE, pchD, pchA* | -1.2 to -2.53 |
| *Pyoverdine* | *pvdA, pvdO, opmQ* | -1.58 to -1.77 |
| *Other proteins* | *fpvA, tseF, PA2384* | -1.34 to -3.51 |
| <u>Antibacterial substances</u> |  |  |
| *Pyocyanin* | *phzA2, phzD2, phzE2, phzF2, phzB1, phzD1, phzE1, phzF1* | -1.32 to -1.70 |
| <u>Virulence factors</u> |  |  |
| *Type III secretion system (T3SS)* | *pscN, popN, pcr1, pcr4, pcrD, pcrG, exsC, pscB, pscE* | 1.20 to 3.06 |
| *Type VI secretion system (T6SS)* |  |  |
| *H1-T6SS* | *tssB1, tssC1, hcp1, tssE1, tssG1, clpV1, vgrG1, tse1, tse3* | 1.22 to 2.02 |
| *H2-T6SS* | *hsiB2, hsiC2, hsiF2, hsiG2, clpV2, sfa2, fha2, lip2, hsiJ2, dotU2, icmF2, pldA* | -1.24 to -1.85 |
| <u>EPS component</u> |  |  |
| *Alginate* | *amrZ* | -1.37 |
| <u>Multidrug resistance</u> |  |  |
| *Drug resistance* | *PA1435, PA3523* | -1.25, -1.44 |
| <u>Two-component system</u> |  |  |
| *Histidine kinase* | *PA0172, ercS', ercS* | -1.31, -1.24, -1.64, |
| *Histidine kinase* | *PA2137* | 2.86 |
| *Response regulator* | *PA0179, gltR* | -1.22, -1.97 |
| *Response regulator* | *PA0756* | 1.81 |
| c-di-GMP |  |  |
| *Diguanylate cyclase* | *siaD, sadC, gcbA, PA4929* | -1.42 to -4.40 |
| *Phosphodiesterase* | *PA2567* | -1.53 |
| *Diguanylate cyclase /phosphodiesterase* | *mucR* | -1.50 |
| Transcription regulators |  |  |
| *Regulators* | *PA0123, PA1196, PA1467, PA1663, PA1864, PA2056, PA2096, ptxS, PA2879, PA3508, PA3714, nalC, PA4596, PA4989* | -1.23 to -3.43 |
| *Regulators* | *PA0217, PA1380, PA1884, PA2334, PA2766, PA3067, glmR* | 1.25 to 2.74 |
| TCA cycle |  |  |
| *Glyoxylate and dicarboxylate metabolism* | *PA0794* | -2.06 |
| *Pyruvate metabolism* | *acoB, PA5445, PA0794* | 1.36, -1.55, -2.06 |
| *Dehydrogenase* | *PA0746, PA2217, PA2552, zwf, fdhA, PA4189, mmsB, mmsA* | -1.40 to -2.78 |

\* Detailed information is provided in the Supplementary Table S3.

Since transcriptome analysis strongly suggested the regulatory role of FleR in bacterial virulence, we next selectively examined productions of two major virulence factors, i.e. pyocyanin and elastase, in *P. aeruginosa*. Our results showed that pyocyanin production was only reduced in the *fleR* deletion mutant, whereas no obvious change was observed in the *fleS* deletion mutant (Fig. 5A), suggesting that regulation of pyocyanin production by FleR is independent of FleS. Interestingly, deletion of either *fleS* or *fleR* resulted in increased elastase production (Fig. 5B), suggesting that elastase in PAO1 is negatively regulated by FleS/FleR. *In trans* expression of wild-type *fleS* and *fleR* in the corresponding *fleS* and *fleR* deletion mutants restored pyocyanin and elastase productions to the wild-type levels (Fig. 5). Together, these results expanded our understanding on the regulatory spectrum of FleS/FleR and confirmed our hypothesis that the TCS FleS/FleR is involved in the regulation of different virulence traits in addition to motility and biofilm formation.

**Fig. 5.**
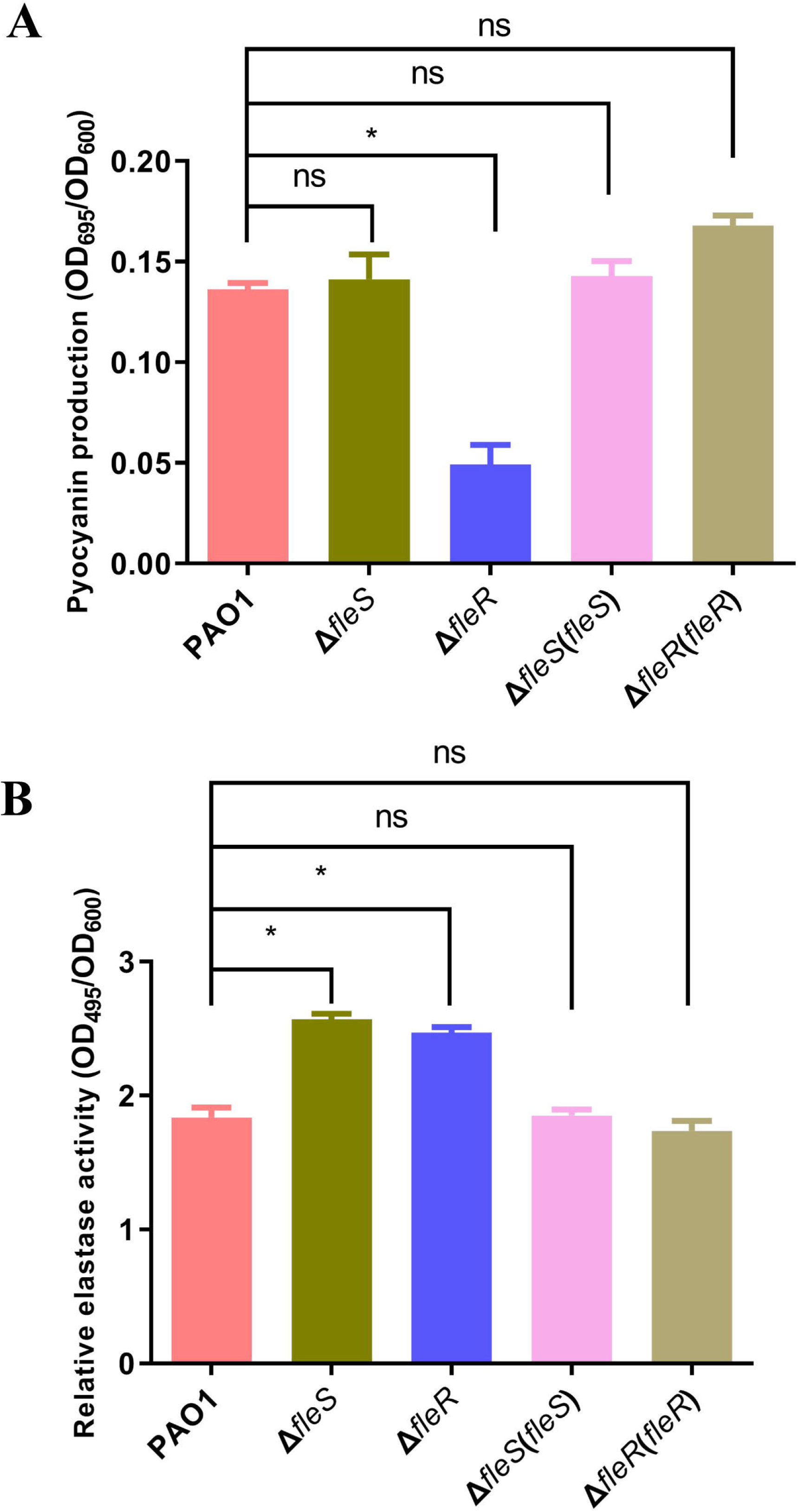
Effects of *fleS* and *fleR* on the elastase and pyocyanin productions. (**A**) Elastase and (**B**) pyocyanin productions in PAO1 and its *fleS* and *fleR* mutants as well as their corresponding complemented strains. The data is the mean of triplicates with standard deviations. *: *p*< 0.05, ns: not significant, tested by Student’s *t*-test.

### FleS and FleR influence the SDS-induced cell aggregation

It has been reported that toxic chemicals, such as antibiotics and detergents, can trigger formation of bacterial cell aggregation which is a bacterial stress response mechanism because aggregated cells are more resistant to biocides than planktonic cells (Drenkard et al., 2003; Gotoh et al., 2008). This phenomenon was also reported in *P. aeruginosa* when it grown in the presence of the detergent SDS (Klebensberger et al., 2006 and 2007). It was shown that aggregated cells exhibited higher survival rate after exposure to SDS than the planktonic cells and the *siaA-D* operon was essential for the SDS-induced formation of cell aggregation (Klebensberger et al., 2007 and 2009). Interestingly, our RNA-seq result and qRT-PCR verification showed that the expression of *siaA-D* genes was significant reduced in the mutant Δ*fleR* (Fig. 6A, Table S3), suggesting that the TCS FleS/FleR is involved in the adaptation to SDS stress. We next moved to examine whether FleS/FleR mediates SDS-induced cell aggregation and found that deletion of either *fleS* or *fleR* indeed substantially reduced formation of cell aggregation (Fig. 6B). In addition, the corresponding complemented strains displayed the fully restored phenotype of cell aggregation, confirming that the TCS FleS/FleR mediates the SDS-induced cell aggregation in PAO1.

**Fig. 6.**
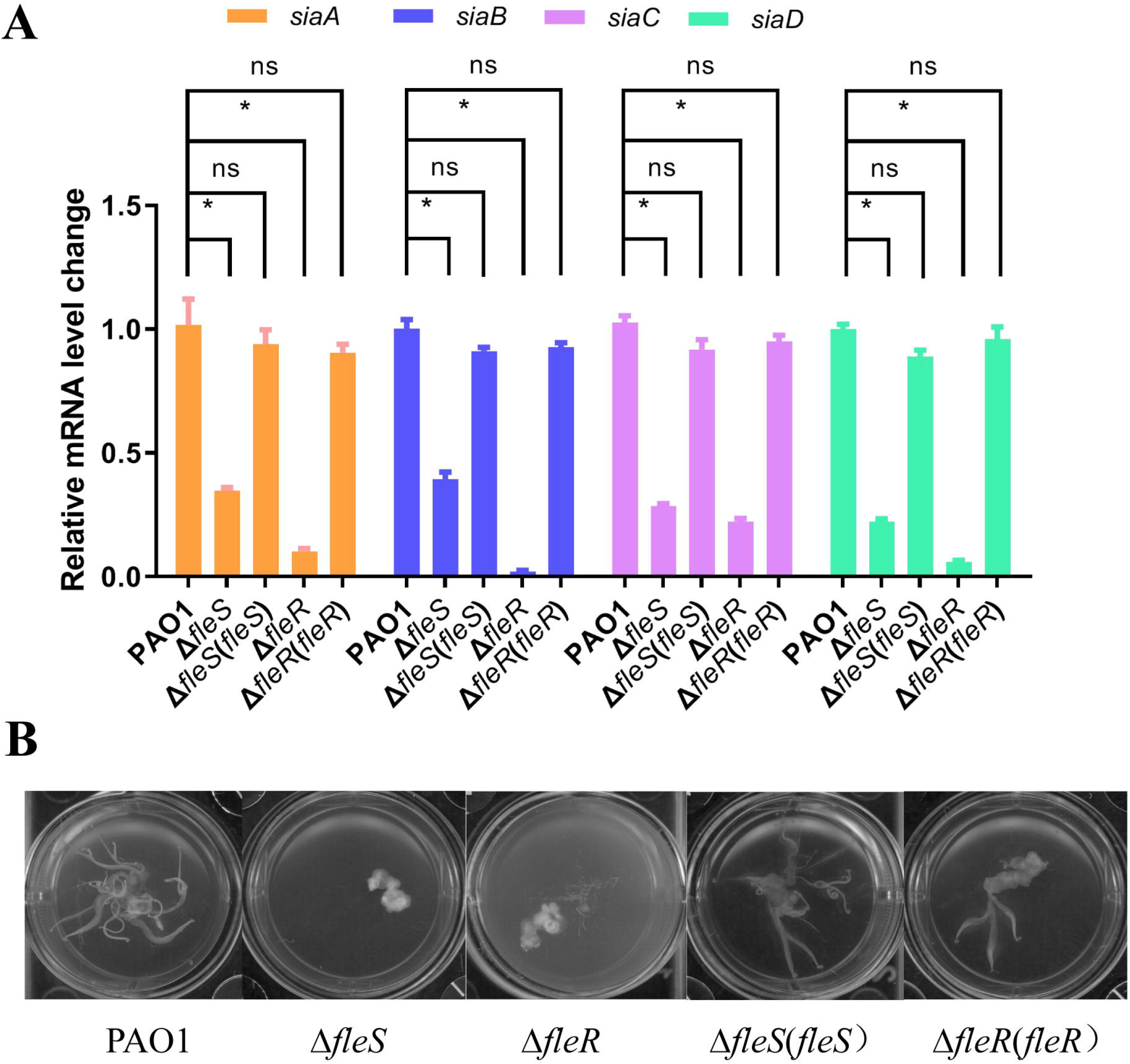
FleS and FleR regulate cell aggregation and *siaA/B/C/D* genes. (**A**) qRT-PCR analysis of the expression of *siaA/B/C/D* genes in the PAO1 strain and its *fleS* and *fleR* derivatives. The *rplU* gene encoding 50S ribosomal protein serves as an internal control. The data is the mean of three replicates with standard deviations. *: *p*< 0.05, ns: not significant, tested by Student’s *t*-test. (**B**) Cell aggregation phenotypes of PAO1 and its derivatives grew in liquid M9 medium containing 0.1% SDS.

### FleR autoactivates the expression of fleSR

Some TCSs are found to autoregulate their own expression such as MisR/MisS in *Neisseria meningitidis* (Tzeng et al., 2006), PhoP/PhoQ in *Salmonella typhimurium* (Newcombe et al., 2004), and CpxR/CpxA in *Escherichia coli* (De et al., 1999). To understand whether FleS/FleR also autoregulates its own expression, we then examined the expression of *fleS* and *fleR* in the mutants Δ*fleR* and Δ*fleS*, respectively. Interestingly, the expression of *fleR* was significantly increased in the *fleS* deletion mutant while the expression of *fleS* was significantly reduced in the *fleR* deletion mutant (Fig. 7A), suggesting that FleR can autoregulate the transcriptional expression of itself, i.e. *fleSR* operon. FleS/FleR autoregulation was further validated by the EMSA assay which examines the binding between FleR and the promoter of *fleSR* (308-bp upstream region of the *fleS* start codon, Fig. 7B). As shown in Fig. 7C, the *fleSR* promoter DNA (P*fleSR*) formed a stable DNA-FleR complex with FleR which migrated at a slower rate than the free probes. Unlabeled probe added in the reaction mix could competitively reduce the amount of labeled DNA in the DNA-FleR complex (Fig. 7C), confirming the specific interaction between the *fleSR* promoter and FleR. These results demonstrated that FleR could autoactivate *fleSR* transcription.

**Fig. 7.**
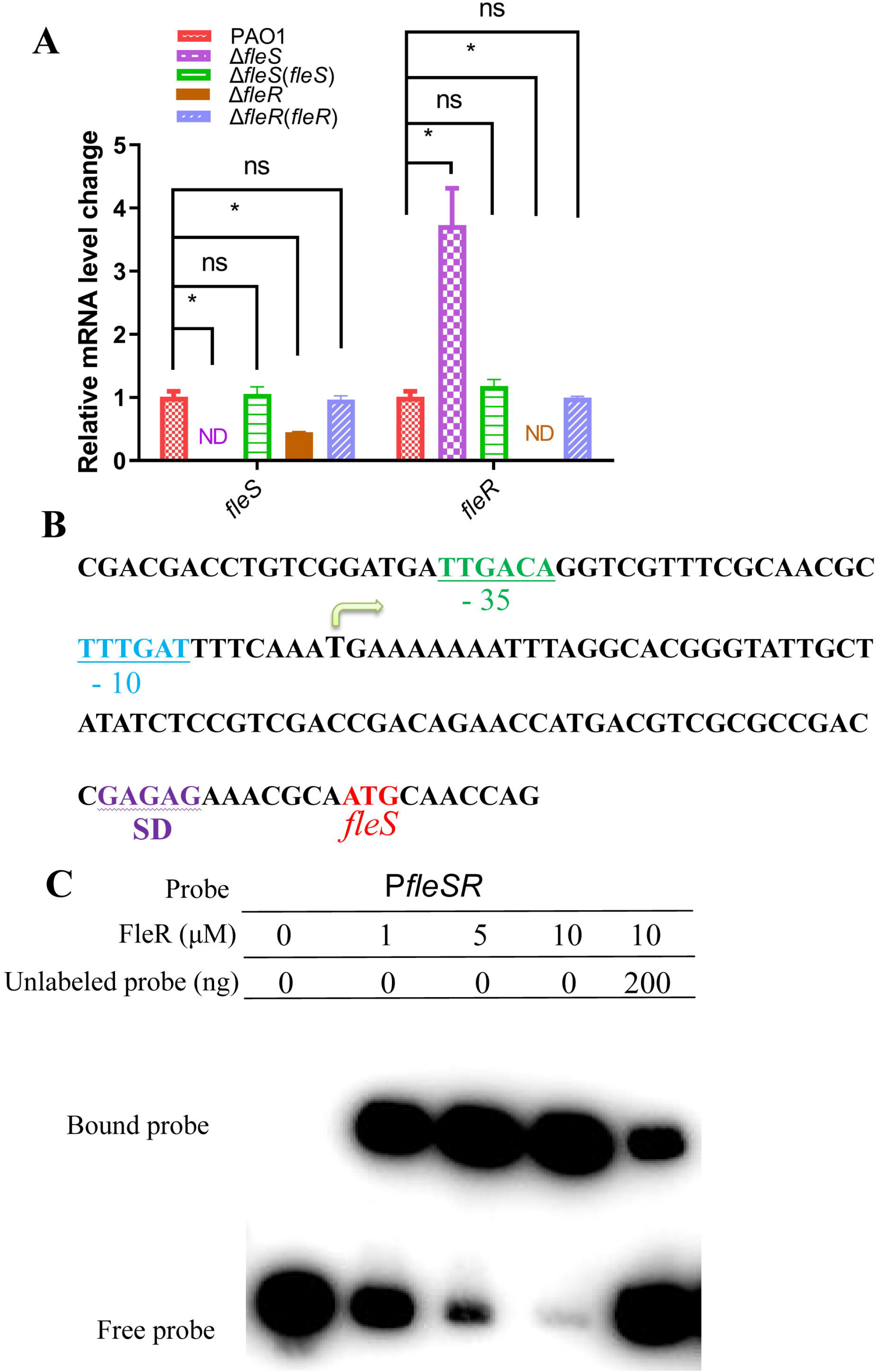
FleR auto-regulates the transcription of *fleSR*. (**A**) qRT-PCR analysis of the expression levels of *fleS* and *fleR* in the *fleS* and *fleR* deletion mutants. The data is the mean of triplicate with standard deviations. *: *p*< 0.05, ns: not significant, tested by Student’s *t*-test. (**B**) Predicted *fleSR* promoter region (P*fleSR*). The putative −10 and −35 elements, the Shine-Dalgarno (SD) sequence, and the *fleS* translation start codon ATG are indicated. (**C**) EMSA examination for the interaction of FleR with P*fleSR*. A biotin-labeled DNA fragment of P*fleSR* is examined with increasing amounts of FleR. For competition analysis, unlabeled P*fleSR* probe is added to the reactions as indicated in the figure.

### FleR regulates flagellum gene expression by direct and specific binding to their promoters

Since our RNA-seq result has suggested that FleR influences bacterial motility by modulating its flagellum biosynthesis, we next sought to understand how FleR regulates the expression of flagellum biosynthesis genes in PAO1. Flagellum biogenesis is known to be controlled by a series of genes in a four-tiered transcriptional regulatory circuit, i.e. *fleQ* in class I, *fleSR*, *flhFfleN*, *fliEFGHIJ*, *fliLMNOPQR, flhB*, *flhA* and *flgA* in class II, *flgBCDE*, *flgFGHIJKL* and *fliK* in class III, and *fliC, fleL*, *cheAB, motAB, cheW*, *cheVR*, *flgMN* and *cheYZ* in class VI (Dasgupta et al., 2003). According to our RNA-seq and qRT-PCR results, the expression of class III and IV genes including *flgBCDE*, *flgFGHIJKL*, *fliK*, *flgMN* and *fliC* was reduced by the absence of *fleR* (Table 1, Table S3, Fig. S3B). We thus conducted EMSA assays to examine whether FleR potentially binds to the promoters of class III and IV genes. The results displayed that the promoters of *flgBCDE*, *flgFGHIJKL and fliC* from the above two classes could form stable complexes with FleR, and the detected interaction signals of FleR with these labeled promoters were enhanced along with the increased levels of FleR and reduced with unlabeled promoters (Fig. 8A, 8B). However, the promoters of *fliK* and *flgMN*, which belong to class III and class IV, respectively, could not form complex with FleR (Fig 8A). Together, these results indicated that FleR directly controls the transcription of *flgBCDE*, *flgFGHIJKL* and *fliC* by interacting with their promoters, and might indirectly modulate the transcription of *flgMN* and *fliK* through other transcription factors.

**Fig. 8.**
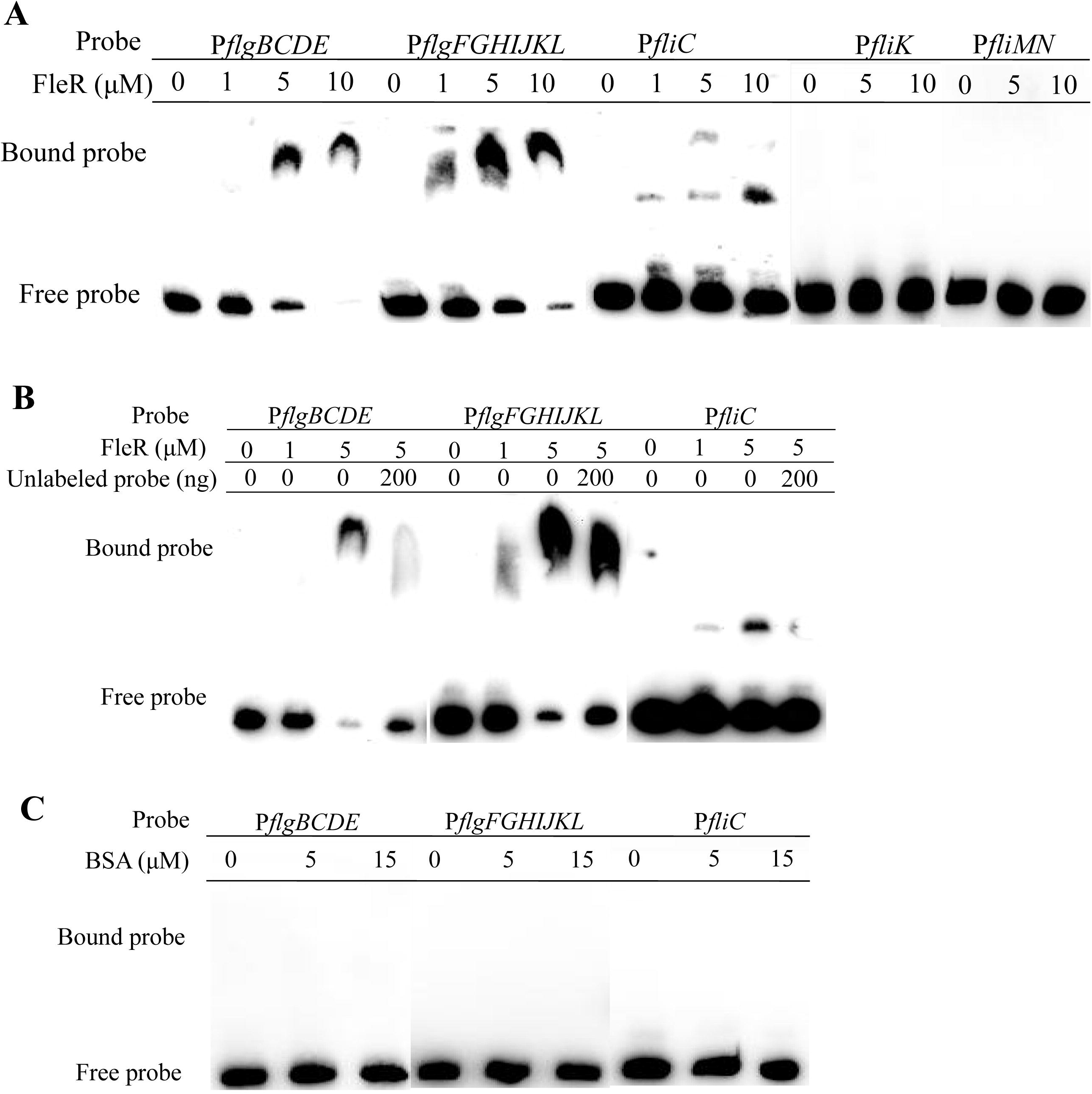
EMSA examinations of FleR binding to the promoter regions of selected flagellum biosynthesis genes. (**A**) EMSA examinations of FleR binding to the promoters of *flgBCDE* (P*flgBCDE*), *flgFGHIJKL* (P*flgFGHIJKL*), *fliC* (P*fliC*), *fliK* (P*flgK*) and *flgMN* (P*flgMN*) with increasing concentrations of FleR. FleR does not bind to P*flgMN* and P*flgK*. (**B**) Analysis of the specific binding of FleR to P*flgBCDE*, P*flgFGHIJKL* and P*fliC* using unlabeled competitive probes which are added to the reaction mixture as indicated. (**C**) Bovine serum albumin (BSA) serves as a negative control which does not interact with P*flgBCDE*, P*flgFGHIJKL* and P*fliC*.

We also performed EMSA analysis to examine the potential interaction of FleR with the promoters of other virulence-related genes such as genes involved in pyocyanin biosynthesis and biofilm formation which are modulated by FleR. Two pyocyanin biosynthesis operons *phz1* and *phz2*, and two biofilm components Pel and Psl exopolysaccharides biosynthesis operons *pel* and *psl* were selected and subjected to EMAS analysis. However, no interaction between FleR and these promoters was observed (Fig. S4), suggesting that FleR might regulate pyocyanin production through modulation of other transcription factor(s) and control the biosynthesis of other component(s) necessary for biofilm formation.

## DISCUSSION

Previous studies reported FleS/FleR might constitute a TCS and suggested its role in the control of bacterial motility through modulating flagellum biogenesis in *P. aeruginosa* (Ritchings et al., 1995; Dasgupta et al., 2003; Kollaran et al., 2019). Recently, *fleS* and *fleR* were also reported to regulate bacterial virulence (Gellatly et al., 2018). However, the regulatory spectrum and the biological functions of FleS/FleR are still largely unknown. In this study, we first showed that FleS and FleR are important to regulate biofilm formation and motility in PAO1 and validated that FleS and FleR constitute a TCS pair (Fig 1, 2 and 3). RNA-seq analysis identified that the expression level of over 400 genes were significantly altered including most of them are virulence-related genes involved in siderophore biosynthesis, pyocyanin biosynthesis, type III/VI secretion systems, c-di-GMP metabolism, flagellar assembly *etc* (Table 1 and S3, Fig. S2, S3). Moreover, FleR was demonstrated to be essential to mediate stress response to SDS (Fig. 6). Lastly, we showed that FleR could directly autoregulate the expression of *fleSR* operon and control bacterial motility by regulating some class III and class IV flagellum biosynthesis genes through directly binding to their promoters (Fig. 7, 8). These findings largely enriched our understanding on the regulatory spectrum and biological roles of this important TCS in *P. aeruginosa*.

It is known that the single polar flagellum of *P. aeruginosa* plays an important role in the bacterial virulence and colonization (Montie et al., 1982; Fleiszig et al., 2001). A previous report revealed the biogenesis of flagellum in this opportunistic pathogen is governed by a four-tiered (Classes I-IV) hierarchy of transcriptional regulation (Dasgupta et al., 2003). Specifically, class I genes are constitutively expressed and include the genes encoding the transcriptional regulator FleQ and the alternative sigma factor FliA (σ^28^). Class II genes include those encoding TCSs such as FleS/FleR which require FleQ and RpoN (σ^54^) for their activation (Jyot et al., 2002). Class III genes are known to be positively regulated by the response regulator FleR in concert with RpoN, and the class IV gene *fliC* is controlled by FliA (Dasgupta et al., 2003). However, the detailed molecular mechanisms of FleR therein remains unclear. Our RNA-seq results showed that deletion of *fleR* significantly decreases the transcription levels of *flgBCDE*, *flgFGHIJKL*, *fliC*, *flgMN* and *fliK* which belong to the III and IV classes, respectively (Table 1, Fig. S3B). Subsequent EMSA analysis demonstrated that FleR regulates the expression of *flgBCDE*, *flgFGHIJKL* and *fliC* by directly binding to their promoters while regulates the expression of *flgMN* and *fliK* indirectly (Fig. 8). Therefore, our results not only provided the molecular evidence that how FleR modulates the expression of class III genes *flgBCDE* and *flgFGHIJKL*, but also revealed that FleR can directly control the expression of the class IV gene *fliC* and indirectly control the expression of additional flagellum synthesis genes, highlighting the versatile roles of FleS/FleR in the regulation of flagellum biogenesis.

qRT-PCR and EMSA analysis indicated that FleR could bind to the promoter region of *fleSR* to activate its own expression (Fig. 7). Although autoregulation of TCS gene expression is found in several TCSs and their regulators have also been shown to bind to their cognate promoter DNA in their non-phosphorylated states such as AgrR in *Cupriavidus metallidurans* and the orphan response regulator in *Streptomyces coelicolor* (Roy et al., 1991; Holman et al., 1994; Liu et al., 1997; Hayde et al., 2002; Wang et al., 2009; Ali et al., 2020), gene transcription without the presence of their cognate sensor kinase is still rare. Interestingly, our study clearly showed that FleR plays a critical role in the activation of its own expression even in the absence of its kinase pair FleS. However, whether FleR can activate gene transcription in the non-phosphorylated state or it could be phosphorylated by other kinase requires further investigation. This is also a possible reason for the discordance in regulation of swimming motility and pyocyanin production by FleS and FleR. Given that FleR shares 39% identity with FleQ whose activity is affected by binding with c-di-GMP and interacting with FleN (Baraquet et al., 2013; Matsuyama et al., 2016), another reason for the activation of FleR without FleS is that FleR might use c-di-GMP or other signaling molecules or interact with other proteins as its activators. Moreover, when we looked into the consensus of promoter sequences of *flgBCDE*, *flgFGHIJKL*, *fliC* and *fleSR* which can be recognized by FleR, no conserved sequence motif among these promoters was found (Fig. S5), suggesting that FleR may have different binding sites in the target gene promoter region. These findings also led us to speculate that the regulatory capacity of FleR requires cooperation with other transcriptional factors such as FleQ, RpoN and PilS/PilR (Jyot et al., 2002; Kilmury et al., 2018).

Over the past three decades, many TCSs from different microorganisms have been functionally characterized with elucidated mechanisms of phosphorylation and regulatory networks. However, identification of the environmental cues which activate various TCSs remains a challenge. Our RNA-seq result identified that the expression of *siaA-D,* which are essential for the SDS-induced formation of cell aggregation, were significantly reduced in the mutant Δ*fleR*, suggesting that the TCS FleS/FleR may be associated with the response to SDS stress. Further analysis validated this speculation as deletion of either *fleS* or *fleR* resulted in reduced formation of bacterial cell aggregates. PAS domain is known to sense diverse intracellular and extracellular signals (Martinez-Argudo et al., 2001; Deng et al., 2012). For example, in *Burkholderia cenocepacia*, the quorum sensing signal BDSF binds to the PAS domain of RpfR with high affinity and activate its phosphodiesterase activity through induction of conformational changes (Deng et al., 2012; Waldron et al., 2019). It is interesting to note that FleS of PAO1 also contains a PAS domain but lacks transmembrane domain, so, it is possible that FleS may perceive intracellular SDS molecules to mediate cell aggregation. However, structural evidence to show the direct interaction of FleS-PAS with SDS is still required.

## CONCLUSIONS

In summary, this study presented a systematic investigation on the regulation and functional roles of the TCS FleS/FleR in *P. aeruginosa*. RNA-seq and phenotype analyses showed that FleS/FleR modulates multiple physiological pathways, including biofilm formation, motility, production of virulence factors, and SDS responsive cell aggregation, highlighting its essential roles in bacterial pathogenicity and adaptation. The molecular mechanisms of FleR in regulating flagellum biosynthesis genes and its own *fleSR* operon were demonstrated. These findings largely enriched our understanding on the spectrum and mechanisms of FleS/FleR in the regulation of bacterial physiology and virulence.

## AUTHOR CONTRIBUTIONS

LZ and TZ designed the experiment. TZ, JH, QF, ZL and QL performed the experiment. TZ analyzed the data. TZ, ZX, and LZ wrote the manuscript.

## Supplementary Table and Figure captions

**Table S1.** Bacterial strains and plasmids used in this study.

**Table S2.** PCR primers used in this study.

**Table S3.** Full list of differentially expressed genes (Log_2_fold change ≥ 1.2) in the *fleR* mutant compared to the wild-type strain. Significantly differentially expressed genes are determined by Cufflinks after Benjamini-Hochberg correction. The fold change is the ratio of the mutant FPKM to the wild-type FPKM.

**Fig. S1.** Growth of PAO1 and its *fleS* and *fleR* mutants as well as corresponding complemented strains. The data is the mean of five replicates with standard deviations.

**Fig. S2.** The relative mRNA levels of the genes involved in pyocyanin biosynthesis (*phzMS*, *phzA1-G1*) (A), c-di-GMP metabolism (*gcbA*, *sadC*, *mucR*, *PA2657*, *PA4929*) and secretion systems (*hcp1*, *tssB1*, *exsC*) (B) in PAO1 and Δ*fleR*. The data is the mean of triplicates with standard deviations. *: *p*< 0.05, ns: not significant, tested by Student’s *t*-test.

**Fig. S3.** The relative mRNA levels of the genes encoding pyoverdine (A) (*pvdA, pvdQ, pvdO, pvdF*), pyochelin (*pchABCD*) and flagellum biosynthesis (B) (*flgB-I*, *fliC*, *flgK*, *flgMN*) in PAO1 and Δ*fleR*. The data is the mean of triplicates with standard deviations. *: *p*< 0.05, ns: not significant, tested by Student’s *t*-test.

**Fig. S4.** EMSA examinations of FleR binding to the promoters of pyocyanin biosynthesis operons *phzA1-G1* (P*phzA1-G1*) and *phzA2-G2* (P*phzA2-G2*), Pel biosynthesis operon *pelA-G* (P*pelA-G*) and Psl biosynthesis operon *pslA-O* (P*pslA-O*).

**Fig. S5.** The promoters of *flgBCDE*, *flgFGHIJKL*, *fliC* and *fleSR* are aligned using clustal X. Blue color represents 100 % identity and pink color represents 75 % identity.

