## Supplemental files for "A Two-Component System FleS/FleR Regulates Multiple Virulence-Related Traits in *Pseudomonas aeruginosa*"

### Supplementary Material

**Table S1.** Bacterial strains and plasmids used in this study.

| Strain or plasmid | Relevant genotype or phenotype | Source or reference |
|---|---|---|
| ***P. aeruginosa*** | | |
| PAO1 | Prototrophic laboratory strain | Lab collection |
| Δ*fleS* | *fleS* deletion mutant of PAO1 with 1208-nt internal coding region deleted | This study |
| Δ*fleR* | *fleR* deletion mutant of PAO1 with 1422-nt internal coding region deleted | This study |
| Δ*fleS*Δ*fleR* | Δ*fleS*Δ*fleR* double-deletion mutant | This study |
| ΔfleS(*fleS*) | Mutant Δ*fleS* harboring the expression construct pBBR1-MCS5-*fleS* | This study |
| Δ*fleR*(*fleR*) | Mutant Δ*fleR* harboring the expression construct pBBR1-MCS5-*fleR* | This study |
| Δ*fleS*Δ*fleR*(*fleS*) | Double-deletion mutant Δ*fleS*Δ*fleR* harboring the expression construct pBBR1-MCS5-*fleS* | This study |
| Δ*fleS*Δ*fleR*(*fleR*) | Double-deletion mutant Δ*fleS*Δ*fleR* harboring the expression construct pBBR1-MCS5-*fleR* | This study |
| ***E. coli* strains** | | |
| DH5α | *spuE44 ΔlacU169(φ80lacZΔM15) hsdR17λpir recA1 endA1 gyrA96 thi-1 relA1* | Lab collection |
| pRK2013 | $Tra^+$, $Mob^-$, ColE1-replicon, $Kan^r$ | Lab collection |
| BL21 | $F^-$*ompT hsdS* ($r_B^-m_B^-$) *dcm*$^+$ $Tet^r$ *gal* (DE3) *endA* | Lab collection |
| **Plasmids** | | |
| pBBR1-MCS5 | Broad host-range cloning vector; $Gm^r$ | Lab collection |
| pGEX-6p-1 | Expression vector, $Amp^r$ | Lab collection |
| pGEX-FleR | pGEX-6p-1 containing *fleR* | This study |
| pBBR1-MCS5-*fleS* | pBBR1-MCS5 containing *fleS* under control of $P_{lac}$ | This study |
| pBBR1-MCS5-*fleR* | pBBR1-MCS5 containing *fleR* under control of $P_{lac}$ | This study |
| pK18mobsacB | Broad-host-range gene replacement vector, sacB,$Gm^r$ | Lab collection |
| pK18-*fleS* | pK18 containing *fleS* flanking regions for generation of *fleS* in-frame deletion | This study |

| | | |
|---|---|---|
| pK18-*fleR* | pK18 containing *fleR* flanking regions for generation of *fleR* in-frame deletion | This study |

Symbol: Gm$^r$, gentamicin resistance; Kan$^r$, kanamycin resistance; Amp$^r$, ampicillin resistance

**Table S2.** PCR primers used in this study.

| Primers | Sequence (5'-3') | Application |
|---|---|---|
| *fleS*-Up-F | gagctcggtacccggggatccGGAGCGCCTGGCGATCAT | For amplification of the 5'-region of *fleS* |
| *fleS*-Up-R | aatcgcagaaagGGCGTTGAGGGCTGGTTG | |
| *fleS*-Dn-F | tcaacgccCTTTCTGCGATTCAGGAGTAACC | For amplification of the 3'-region of *fleS* |
| *fleS*-Dn-R | acgacggccagtgccaagcttCGCCGGAGATCAGCACGG | |
| *fleR*-Up-F | gagctcggtacccggggatccGCTGGTGTTCGCCCGCGG | For amplification of the 5'-region of *fleR* |
| *fleR*-Up-R | agcacGGGGTTACTCCTGAATCGCAG | |
| *fleR*-Dn-F | ttcaggagtaaccccGTGCTCGCCATGTTCCCC | For amplification of the 3'-region of *fleR* |
| *fleR*-Dn-R | acgacggccagtgccaagcttACGCTGGCCTTCTGGCTG | |
| *fleS*-FC-F | gtcgacggtatcgataagcttGGGGTGATCGGGGTCGGC | For construction of pBBR1-MCS5-*fleS* |
| *fleS*-FC-R | cgctctagaactagtggatccCGCGTAGTGCGCGGTCGT | |
| *fleR*-FC-F | gtcgacggtatcgataagcttTGCGGGCCCGAACTGCGC | For construction of pBBR1-MCS5-*fleR* |
| *fleR*-FC-R | cgctctagaactagtggatccGCGGACGCAAAAGGCCCG | |
| FleR-GST-F | ttccaggggcccctgggatccATGGCAGCCAAAGTCCTGC | For expression of FleR-GST fusion protein |
| FleR-GST-R | ctcgagtcgacccgggaattcTCAGATGGCGTAGAGATAGGCC | |
| *fleSR*operon-F | CACCAGGGGCAATTGCAGTTG | |
| *fleSR*operon-R1 | GTGTCGCTGAGGGCTTCGCGTA | |
| *fleSR*operon-R2 | CCGTCCATGCCCGGCATGTTCAC | For RT-PCR analysis |
| *fleSR*operon-R3 | GTAGGCGGTCATCAGCAACAC | |

| | | |
|---|---|---|
| EMSA-*flgBCDE*-F | GCGATACCGCCTTCAACTTC | |
| EMSA-*flgBCDE*-R | CTGCTGGTGGATACCGAGTG | |
| EMSA-*flgFGHIJKL*-F | CGATCGGCAAGACCTCCT | For EMSA analysis of FleR |
| EMSA-*flgFGHIJKL*-R | TTGGCATGAGCACGCATC | |
| EMSA-*fliC*-F | ACGGGAGGGCTAAAGAAAATCG | |
| EMSA-*fliC*-R | TGTTCGTGTTGACTGTAAGGGCC | |
| EMSA-*fleS*-F | AAGGCCTGGACCTCAAGGAC | |
| EMSA-*fleS*-R | GCTGGTTGCATTGCGTTTC | |
| q-*flgB*-F | GAGCACTCGGTATCCACCAG | |
| q-*flgB*-R | GATATGCCGCTCGTTGGTG | |
| q-*flgC*-F | CCTCGCCAGTGTCTTCAACA | |
| q-*flgC*-R | CTGCTGGAACATGGTGGAGA | |
| q-*flgD*-F | GTCGCTGAACAAGAGCATGG | |
| q-*flgD*-R | TTGACCCATACGTTGCTGCT | |
| q-*flgF*-F | GTGCTCATGCCAACAACCTG | |
| q-*flgF*-R | GCCATGGCTGAAATCGGT | |
| q-*flgE*-F | ACCTCAACTCGACGCTGAAG | |
| q-*flgE*-R | AGAACTGGCTCATGGTGTGG | |
| q-*flgG*-F | TCAACGTGGTCGAGGAACTG | |
| q-*flgG*-R | CTGGGTGACGAAGGACAACA | |
| q-*flgH*-F | CAGAACCTCTACGACGACCG | |
| q-*flgH*-R | TCGGAGTTGGCTTTCTTGCT | |
| q-*flgI*-F | ACCAGTTGATCGGCTATGGC | |
| q-*flgI*-R | TGTTGAAGGTCTGCAGGGTG | |
| q-*flgK*-F | GCCGGTTGGAGTTATGTCGA | |
| q-*flgK*-R | GCACGTTGGACAGGTTGAAC | For qRT-PCR analysis |
| q-*flgM*-F | TGCAGAAAGTCACGGACCAG | |
| q-*flgM*-R | TCGAAGTCGAGCAGCTTGC | |
| q-*flgN*-F | TGCAGTTGGTCGACGAAGAA | |
| q-*flgN*-R | TCCAGTTGCTGCATCAGTGG | |
| q-*fliC*-F | CCTGCAGCAGTCCACCAATA | |
| q-*fliC*-R | CGGAGATACGGGTCAGTTCG | |
| q-*hcp1*-F | TCCAAGGACAAGACTCACGC | |
| q-*hcp1*-R | TTGGTGAACGACAGGTCCTG | |
| q-*tssB1*-F | AGAGCATGGACGACTTCAGC | |
| q-*tssB1*-R | ATGTAGGTCAGCAGGTTGGC | |
| q-*exsC*-F | CGATCTGCATTTCGGCTTCG | |
| q-*exsC*-R | TGATCGAGCAGATTGGCCAA | |
| q-*phzM*-F | TCGGCGAAGACTTCTACAGC | |
| q-*phzM*-R | GATGGCCTTGGTCAATTCGC | |

| | | |
|---|---|---|
| q-*phzS*-F | GGCGTCGGCATCAATATCCA | |
| q-*phzS*-R | AGTACTGCGGATAGGCGTTG | |
| q-*phzA1*-F | GGATCCGAACCACTTCTGGG | |
| q-*phzA1*-R | GCTATTCCCAATGCACGCAG | |
| q-*phzB1*-F | CACCGATGCCAACGAACTTC | |
| q-*phzB1*-R | GGCATGTTCCGCAAGTTTGT | |
| q-*phzC1*-F | GATCCTCAAGGGCTATGCGG | |
| q-*phzC1*-R | GTCGAACCGAGATAGACCCG | |
| q-*phzD1*-F | GACATGCAGCGCTACTTCCT | |
| q-*phzD1*-R | AGAAGTCCTTGAGCAGACCG | |
| q-*phzE1*-F | GATTCCCTACCGGCAGATCG | |
| q-*phzE1*-R | CGAAACCTTCCTCGCTCAGT | |
| q-*phzF1*-F | CGACAAGGACAGGCTGTTCC | |
| q-*phzF1*-R | CCTCGATGGGAAAGGTCGAG | |
| q-*phzG1*-F | TACCAGACCCACCCGATGTC | |
| q-*phzG1*-R | GACTCCAGGCACAGTTCGAA | |
| q-*pchA*-F | GAACGCTACCGGGTACTCTG | |
| q-*pchA*-R | GCTCACCTTGGCTTCCCATT | |
| q-*pchB*-F | CCGGATCGACCTGGATATCG | |
| q-*pchB*-R | GAACAGTCCCTCGACGAAGG | |
| q-*pchC*-F | CTGTTCGTCTCCGCCCATC | For qRT-PCR analysis |
| q-*pchC*-R | GTCTCGATCGCCTGGTAGTC | |
| q-*pvdA*-F | GGAAGTGCTGTTCCTGGACA | |
| q-*pvdA*-R | GCCCAGGTTGATGAAGTCGA | |
| q-*pvdO*-F | AAGAAGACCGGCAAACGCTA | |
| q-*pvdO*-R | GCGAGGTGAAGTTGTAGCCA | |
| q-*pvdQ*-F | GGACTTCGTGCAGAACTCCA | |
| q-*pvdQ*-R | CCATCTCCTCGAGCGTCTTC | |
| q-*pvdF*-F | CAGCGGTACATGAAGTCGGT | |
| q-*pvdF*-R | AGGCGAAACCGTAGTCCTTG | |
| q-*fleS*-F | ACGGTCACGAGATTTCCCTG | |
| q-*fleS*-R | GGTATCGGTGAGGTCGTTGAG | |
| q-*fleR*-F | GCCTGATCCGTACACGCTAC | |
| q-*fleR*-R | GAACGGCTTGACCAGGTAGTC | |
| *rplU*-F | GCAGCACAAAGTCACCGAAG | Internal control for |
| *rplU*-R | CCGATTTTCACGTCTTCGCC | qRT-PCR analysis |

**Table S3.** Full list of differentially expressed genes ($Log_2$fold change ≥ 1.2) in the fleR mutant compared to the wild-type strain. Significantly differentially expressed genes are determined by Cufflinks after Benjamini-Hochberg correction. The fold change is the ratio of the mutant FPKM to the wild-type FPKM.

| Locus tag | Gene name | Fold chang | Gene function | Number of pathway // Name of pathway |
|---|---|---|---|---|
| PA0045 | PA0045 | -1.990 | curli production assembly protein CsgG | - |
| PA0046 | PA0046 | -2.690 | DUF4810 domain-containing protein | - |
| PA0047 | PA0047 | -1.234 | lipoprotein | - |
| PA0075 | pppA | -1.541 | serine/threonine phosphatase | ko03070//Bacterial secretion system |
| PA0083 | PA0083 | 1.227 | type VI secretion system-associated protein | - |
| PA0084 | PA0084 | 1.305 | EvpB/family type VI secretion protein | - |
| PA0085 | hcp1 | 1.285 | protein secretion apparatus assembly protein | ko03070//Bacterial secretion system |
| PA0087 | PA0087 | 1.224 | type VI secretion system lysozyme | - |
| PA0089 | PA0089 | 1.350 | type VI secretion protein | |
| PA0090 | clpV1 | 1.822 | secretion protein ClpV1 | ko03070//Bacterial secretion system |
| PA0091 | vgrG1 | 1.612 | type VI secretion system protein VgrG | ko03070//Bacterial secretion system |
| PA0103 | PA0103 | 2.038 | sulfate transporter | - |
| PA0117 | PA0117 | -1.846 | short-chain dehydrogenase | - |
| PA0119 | PA0119 | -1.876 | C4-dicarboxylate transporter 1 | ko02020//Two-component system |
| PA0122 | PA0122 | -1.296 | hemolysin | - |
| PA0123 | PA0123 | -1.377 | transcriptional regulator | - |
| PA0126 | PA0126 | -1.203 | Uncharacterised protein | - |
| PA0128 | PA0128 | 1.725 | alkylphosphonate utilization protein | - |
| PA0137 | PA0137 | 1.395 | ABC transporter permease | - |
| PA0162 | opdC | -1.464 | histidine porin OpdC | - |
| PA0169 | PA0169 | -2.938 | GGDEF domain-containing protein | - |
| PA0170 | PA0170 | -2.323 | Fe-S oxidoreductase | - |
| PA0171 | PA0171 | -3.016 | Uncharacterized protein PAE221_01065 | - |
| PA0172 | PA0172 | -1.313 | histidine kinase | - |
| PA0179 | PA0179 | -1.227 | two-component response regulator | ko02020//Two-component system;ko02030//Bacterial chemotaxis |

| | | | | |
|---|---|---|---|---|
| PA0195 | pntAA | 1.290 | NAD(P) transhydrogenase subunit alpha | ko00760//Nicotinate and nicotinamide metabolism |
| PA0199 | exbD1 | -1.205 | biopolymer transport protein ExbD | - |
| PA0200 | PA0200 | -1.284 | DUF3079 domain-containing protein | - |
| PA0215 | PA0215 | 1.533 | malonate transporter MadL | - |
| PA0217 | PA0217 | 1.247 | transcriptional regulator | - |
| PA0222 | PA0222 | -1.435 | ABC transporter, periplasmic spermidine putrescine-binding protein PotD | - |
| PA0238 | PA0238 | 2.137 | xylose isomerase | - |
| PA0251 | PA0251 | 1.858 | suppressor of fused family protein | - |
| PA0252 | PA0252 | -1.998 | Uncharacterized protein PAE221_00084 | - |
| PA0263 | hcpC | -1.311 | major exported protein, partial | ko03070//Bacterial secretion system |
| PA0280 | cysA | -2.114 | sulfate.thiosulfate ABC transporter ATP-binding protein CysA | ko02010//ABC transporters;ko00920//Sulfur metabolism |
| PA0281 | cysW | -1.678 | sulfate transporter CysW | ko02010//ABC transporters;ko00920//Sulfur metabolism |
| PA0282 | cysT | -2.519 | sulfate transporter CysT | ko02010//ABC transporters;ko00920//Sulfur metabolism |
| PA0320 | PA0320 | -1.614 | TIGR00156 family protein | - |
| PA0323 | PA0323 | -1.629 | Putrescine-binding periplasmic protein precursor | ko02010//ABC transporters |
| PA0451 | PA0451 | 1.312 | peptidase | - |
| PA0471 | PA0471 | -1.628 | transmembrane sensor | - |
| PA0472 | PA0472 | -1.899 | RNA polymerase sigma factor | - |
| PA0493 | PA0493 | 13.316 | probable biotin-requiring enzyme | - |
| PA0513 | PA0513 | -1.387 | heme d1 biosynthesis protein NirG | - |
| PA0517 | nirC | -1.356 | cytochrome c55X | - |
| PA0519 | nirS | -1.273 | nitrite reductase | ko00910//Nitrogen metabolism |
| PA0524 | norB | -1.310 | nitric oxide reductase subunit B | ko00910//Nitrogen metabolism |

| | | | | |
|---|---|---|---|---|
| PA0534 | PA0534 | -1.867 | Oxidoreductase | - |
| PA0579 | rpsU | 1.431 | 30S ribosomal protein S21 | ko03010//Ribosome |
| PA0672 | hemO | -1.450 | heme oxygenase | - |
| PA0674 | vreA | -4.254 | VreA | - |
| PA0680 | PA0680 | 3.410 | HxcV pseudopilin | ko03070//Bacterial secretion system |
| PA0734 | PA0734 | 1.949 | cysteine-rich CWC family protein | - |
| PA0744 | PA0744 | -1.964 | enoyl-CoA hydratase | - |
| PA0745 | PA0745 | -1.656 | enoyl-CoA hydratase | - |
| PA0746 | PA0746 | -1.396 | acyl-CoA dehydrogenase | ko00281//Geraniol degradation |
| PA0753 | PA0753 | -2.888 | tripartite tricarboxylate transporter TctB family protein | ko02020//Two-component system |
| PA0754 | PA0754 | -1.291 | C4-dicarboxylate ABC transporter substrate-binding protein | ko02020//Two-component system |
| PA0755 | opdH | -1.437 | cis-aconitate porin OpdH | - |
| PA0756 | PA0756 | 1.810 | two-component response regulator | ko02020//Two-component system |
| PA0793 | PA0793 | -1.726 | 3-methylitaconate isomerase | - |
| PA0794 | PA0794 | -2.059 | aconitate hydratase | ko01230//Biosynthesis of amino acids;ko01200//Carbon metabolism;ko00630//Glyoxylate and dicarboxylate metabolism;ko00020//Citrate cycle (TCA cycle);ko01210//2-Oxocarboxylic acid metabolism |
| PA0795 | prpC | -2.380 | methylcitrate synthase | ko00640//Propanoate metabolism |
| PA0796 | prpB | -1.562 | 2-methylisocitrate lyase | ko00640//Propanoate metabolism |
| PA0837 | slyD | 1.240 | peptidyl-prolyl cis-trans isomerase SlyD | - |
| PA0855 | PA0855 | -1.275 | Protein of uncharacterised function (DUF2804) | - |

| | | | | |
|---|---|---|---|---|
| PA0865 | hpd | -1.350 | 4-hydroxyphenylpyruvate dioxygenase | ko00350//Tyrosine metabolism;ko00360//Phenylalanine metabolism;ko00130//Ubiquinone and other terpenoid-quinone biosynthesis |
| PA0866 | aroP2 | -2.383 | aromatic amino acid transporter AroP | - |
| PA0887 | acsA | -3.339 | acetyl-CoA synthetase | ko01200//Carbon metabolism;ko00620//Pyruvate metabolism;ko00640//Propanoate metabolism;ko00010//Glycolysis / Gluconeogenesis;ko00680//Methane metabolism |
| PA0909 | PA0909 | 2.979 | putative membrane protein | - |
| PA0939 | PA0939 | -1.714 | Rho-specific inhibitor of transcription termination YaeO | - |
| PA0960 | PA0960 | 1.278 | slyX protein | - |
| PA1058 | PA1058 | 12.878 | monovalent cation/H+ antiporter subunit F | - |
| PA1070 | braG | -1.288 | branched-chain amino acid ABC transporter ATP-binding protein BraG | ko02010//ABC transporters |
| PA1071 | braF | -1.259 | branched-chain amino acid ABC transporter ATP-binding protein BraF | ko02010//ABC transporters |
| PA1077 | flgB | -4.925 | flagellar basal-body rod protein FlgB | ko02040//Flagellar assembly |
| PA1078 | flgC | -4.174 | flagellar basal body rod protein FlgC | ko02040//Flagellar assembly |
| PA1079 | flgD | -4.029 | flagellar basal body rod modification protein | ko02040//Flagellar assembly |
| PA1080 | flgE | -4.774 | flagellar hook protein FlgE | ko02040//Flagellar assembly |
| PA1081 | flgF | -4.232 | flagellar basal body rod protein FlgF | ko02040//Flagellar assembly |
| PA1092 | fliC | -1.763 | flagellin type B | ko02020//Two-component system;ko02040//Flagellar assembly |
| PA1099 | fleR | -5.866 | two-component response regulator | ko02020//Two-component system |
| PA1132 | PA1132 | -1.441 | hypothetical protein | - |
| PA1137 | PA1137 | -1.860 | oxidoreductase | - |

| | | | | |
|---|---|---|---|---|
| PA1168 | PA1168 | -1.396 | Uncharacterised protein | - |
| PA1196 | PA1196 | -2.201 | transcriptional regulator | - |
| PA1197 | PA1197 | -1.565 | NAD-dependent protein deacylase | - |
| PA1213 | PA1213 | -1.400 | clavaminic acid synthetase | - |
| PA1240 | PA1240 | -1.955 | enoyl-CoA hydratase | - |
| PA1254 | PA1254 | -1.709 | dihydrodipicolinate synthetase | ko01230//Biosynthesis of amino acids;ko00300//Lysine biosynthesis;ko00261//Monobactam biosynthesis |
| PA1255 | PA1255 | -1.545 | trans-3-hydroxy-L-proline dehydratase | - |
| PA1266 | PA1266 | -2.065 | oxidoreductase | - |
| PA1267 | PA1267 | -1.839 | FAD binding domain protein | - |
| PA1275 | cobD | -1.843 | cobalamin biosynthesis protein | ko00860//Porphyrin and chlorophyll metabolism |
| PA1276 | cobC | -1.723 | threonine-phosphate decarboxylase | ko00860//Porphyrin and chlorophyll metabolism |
| PA1277 | cobQ | -1.716 | cobyric acid synthase | ko00860//Porphyrin and chlorophyll metabolism |
| PA1279 | cobU | -1.572 | nicotinate-nucleotide--dimethylbenzimidazole phosphoribosyltransferase | ko00860//Porphyrin and chlorophyll metabolism |
| PA1281 | cobV | -3.316 | adenosylcobinamide-GDP ribazoletransferase | ko00860//Porphyrin and chlorophyll metabolism |
| PA1282 | PA1282 | -1.803 | MFS transporter | - |
| PA1286 | PA1286 | 1.818 | major facilitator superfamily transporter | - |
| PA1319 | cyoC | -1.284 | cytochrome o ubiquinol oxidase subunit III | ko00190//Oxidative phosphorylation |
| PA1343 | PA1343 | -1.235 | sn-glycerol-3-phosphate transporter | - |
| PA1349 | PA1349 | 1.297 | dehydrogenase | - |
| PA1353 | PA1353 | 1.736 | VOC family protein | - |
| PA1380 | PA1380 | 1.365 | transcriptional regulator | - |
| PA1412 | PA1412 | 1.651 | major Facilitator Superfamily protein | - |

| | | | | |
|---|---|---|---|---|
| PA1417 | PA1417 | -1.231 | probable decarboxylase | ko01230//Biosynthesis of amino acids;ko00650//Butanoate metabolism;ko01210//2-Oxocarboxylic acid metabolism;ko00770//Pantothenate and CoA biosynthesis;ko00290//Valine, leucine and isoleucine biosynthesis;ko00660//C5-Branched dibasic acid metabolism |
| PA1420 | PA1420 | 3.131 | Undecaprenyl pyrophosphate synthase | - |
| PA1423 | bdlA | -1.469 | pili assembly chaperone | ko02020//Two-component system;ko02030//Bacterial chemotaxis |
| PA1428 | PA1428 | 1.301 | GNAT family N-acetyltransferase | - |
| PA1435 | PA1435 | -1.252 | resistance-nodulation-cell division (RND) efflux membrane fusion protein | - |
| PA1441 | PA1441 | -2.471 | flagellar hook-length control protein FliK | ko02040//Flagellar assembly |
| PA1442 | PA1442 | 1.282 | flagellar basal body protein FliL | - |
| PA1467 | PA1467 | -2.516 | transcriptional regulator | - |
| PA1476 | ccmB | 1.304 | heme exporter protein B | ko02010//ABC transporters |
| PA1487 | PA1487 | -1.280 | carbohydrate kinase | ko00561//Glycerolipid metabolism |
| PA1498 | pykF | 1.853 | pyruvate kinase | ko01230//Biosynthesis of amino acids;ko01200//Carbon metabolism;ko00230//Purine metabolism;ko00620//Pyruvate metabolism;ko00010//Glycolysis / Gluconeogenesis |
| PA1500 | PA1500 | -1.378 | oxidoreductase | ko00630//Glyoxylate and dicarboxylate metabolism |
| PA1545 | PA1545 | -1.612 | Uncharacterised protein | - |

| | | | | |
|---|---|---|---|---|
| PA1561 | aer | -1.780 | aerotaxis receptor Aer | ko02020//Two-component system;ko02030//Bacterial chemotaxis |
| PA1594 | PA1594 | 1.231 | thioesterase, partial | - |
| PA1596 | htpG | -1.902 | chaperone protein HtpG | - |
| PA1608 | PA1608 | -1.388 | chemotaxis transducer | ko02020//Two-component system;ko02030//Bacterial chemotaxis |
| PA1635 | kdpC | -2.520 | potassium-transporting ATPase subunit C | ko02020//Two-component system |
| PA1638 | PA1638 | 1.235 | glutaminase | ko00250//Alanine, aspartate and glutamate metabolism;ko00220//Arginine biosynthesis;ko00471//D-Glutamine and D-glutamate metabolism |
| PA1657 | PA1657 | -1.517 | type VI secretion system-associated protein | - |
| PA1658 | PA1658 | -1.572 | EvpB/family type VI secretion protein | - |
| PA1659 | PA1659 | -1.241 | type VI secretion protein | - |
| PA1660 | PA1660 | -1.291 | type VI secretion protein | - |
| PA1662 | PA1662 | -1.393 | ClpA/B-type protease | ko03070//Bacterial secretion system |
| PA1663 | PA1663 | -1.560 | transcriptional regulator | - |
| PA1665 | PA1665 | -1.299 | signal peptide protein | - |
| PA1666 | PA1666 | -1.372 | Type VI secretion lipoprotein/VasD | ko03070//Bacterial secretion system |
| PA1667 | PA1667 | -1.634 | type VI secretion protein | - |
| PA1668 | PA1668 | -1.852 | membrane protein | ko03070//Bacterial secretion system |
| PA1669 | PA1669 | -1.546 | type VI secretion protein IcmF | ko03070//Bacterial secretion system |
| PA1671 | stk1 | -1.234 | serine-threonine kinase Stk1 | - |
| PA1679 | PA1679 | -2.013 | Uncharacterized protein PAE221_02692 | - |
| PA1698 | popN | 1.485 | type III secretion outer membrane protein PopN | ko03070//Bacterial secretion system |
| PA1699 | PA1699 | 1.227 | Pcr1 | - |
| PA1702 | PA1702 | 3.059 | chaperone protein yscY | - |
| PA1703 | pcrD | 1.205 | type III secretory apparatus protein PcrD | ko03070//Bacterial secretion system |

| | | | | |
|---|---|---|---|---|
| PA1705 | pcrG | 1.250 | type III secretion regulator | - |
| PA1710 | exsC | 1.224 | exoenzyme S synthesis protein ExsC | - |
| PA1715 | pscB | 1.291 | type III export apparatus protein | - |
| PA1718 | pscE | -1.382 | type III export protein PscE | - |
| PA1727 | mucR | -1.502 | bifunctional diguanylate cyclase/phosphodiesterase | - |
| PA1747 | PA1747 | -1.327 | putative secreted protein | - |
| PA1844 | PA1844 | 2.029 | Uncharacterized protein PAE221_01822 | - |
| PA1845 | PA1845 | 2.353 | Uncharacterised protein | - |
| PA1852 | PA1852 | 1.872 | Uncharacterized protein PAE221_01813 | - |
| PA1864 | PA1864 | -1.775 | transcriptional regulator | - |
| PA1884 | PA1884 | 1.270 | transcriptional regulator | - |
| PA1888 | PA1888 | -1.264 | Uncharacterised protein | - |
| PA1899 | phzA2 | 1.337 | phenazine biosynthesis protein | - |
| PA1902 | phzD2 | -1.698 | phenazine biosynthesis protein PhzD | - |
| PA1903 | phzE2 | -1.324 | phenazine biosynthesis protein PhzE | - |
| PA1904 | phzF2 | -1.369 | 2,3-dihydro-3-hydroxyanthranilate isomerase | - |
| PA1912 | femI | -2.770 | ECF sigma factor FemI | - |
| PA1913 | PA1913 | -5.281 | Uncharacterized protein PAE221_00740 | - |
| PA1920 | nrdD | -1.281 | anaerobic ribonucleoside triphosphate reductase | ko00230//Purine metabolism;ko00240//Pyrimidine metabolism |
| PA1947 | rbsA | -1.334 | ribose transporter RbsA | ko02010//ABC transporters |
| PA1948 | rbsC | -1.340 | ABC transporter permease | ko02010//ABC transporters |
| PA1957 | PA1957 | 1.270 | N-acetylglucosamine-6-sulfatase | - |
| PA1967 | PA1967 | -2.970 | Uncharacterised protein | - |
| PA1968 | PA1968 | 1.207 | Uncharacterised protein | - |
| PA1976 | ercS' | -1.238 | hybrid sensor histidine kinase/response regulator | - |
| PA1988 | pqqD | -1.693 | coenzyme PQQ synthesis protein D | - |

| | | | | |
|---|---|---|---|---|
| PA1992 | ercS | -1.635 | sensor histidine kinase | - |
| PA2000 | dhcB | -1.397 | Succinyl-CoA:3-ketoacid coenzyme A transferase subunit B | ko00280//Valine, leucine and isoleucine degradation;ko00650//Butanoate metabolism;ko00072//Synthesis and degradation of ketone bodies |
| PA2001 | atoB | -1.365 | acetyl-CoA acetyltransferase | ko02020//Two-component system;ko01200//Carbon metabolism;ko00620//Pyruvate metabolism;ko00630//Glyoxylate and dicarboxylate metabolism;ko01212//Fatty acid metabolism;ko00280//Valine, leucine and isoleucine degradation;ko00640//Propanoate metabolism;ko00650//Butanoate metabolism;ko00362//Benzoate degradation;ko00071//Fatty acid degradation;ko00380//Tryptophan metabolism;ko00310//Lysine degradation;ko00900//Terpenoid backbone biosynthesis;ko00072//Synthesis and degradation of ketone bodies |
| PA2004 | PA2004 | -1.326 | citrate transporter family protein | - |
| PA2013 | liuC | -1.311 | gamma-carboxygeranoyl-CoA hydratase | ko00280//Valine, leucine and isoleucine degradation |
| PA2027 | PA2027 | 2.036 | Uncharacterized protein PAE221_00397 | - |
| PA2029 | PA2029 | 1.285 | winged helix-turn helix family protein | - |
| PA2036 | PA2036 | -1.579 | methyltransferase domain protein | - |

| | | | | |
|---|---|---|---|---|
| PA2040 | PA2040 | -1.249 | glutamine synthetase | ko02020//Two-component system;ko01230//Biosynthesis of amino acids;ko00630//Glyoxylate and dicarboxylate metabolism;ko00910//Nitrogen metabolism;ko00250//Alanine, aspartate and glutamate metabolism;ko00220//Arginine biosynthesis |
| PA2048 | PA2048 | -12.152 | antibiotic biosynthesis monooxygenase | - |
| PA2056 | PA2056 | -1.745 | LysR family transcriptional regulator | - |
| PA2061 | PA2061 | 1.617 | ABC transporter ATP-binding protein | ko02010//ABC transporters |
| PA2096 | PA2096 | -1.316 | transcriptional regulator | - |
| PA2099 | PA2099 | -2.643 | short-chain dehydrogenase | - |
| PA2110 | PA2110 | -2.114 | allophanate hydrolase subunit 2 family protein | - |
| PA2112 | PA2112 | -1.807 | LamB/YcsF family protein | - |
| PA2113 | opdO | -1.439 | pyroglutatmate porin | - |
| PA2134 | PA2134 | 1.335 | membrane protein | - |
| PA2136 | PA2136 | -1.688 | Uncharacterized protein PAE221_01703 | - |
| PA2137 | PA2137 | 2.860 | histidine kinase | - |
| PA2138 | PA2138 | 3.330 | multifunctional non-homologous end joining protein LigD | ko03450//Non-homologous end-joining |
| PA2147 | katE | 1.410 | catalase HPII | ko01200//Carbon metabolism;ko00630//Glyoxylate and dicarboxylate metabolism;ko00380//Tryptophan metabolism |
| PA2148 | PA2148 | 1.813 | methyltransferase | - |
| PA2154 | PA2154 | 1.309 | membrane protein | - |
| PA2160 | PA2160 | 1.920 | glycosyl hydrolase | ko00500//Starch and sucrose metabolism |
| PA2161 | PA2161 | 2.983 | Protein of uncharacterised function (DUF2934) | - |

| | | | | |
|---|---|---|---|---|
| PA2162 | PA2162 | 1.438 | malto-oligosyltrehalose synthase | ko00500//Starch and sucrose metabolism |
| PA2164 | PA2164 | 1.268 | glycosyl hydrolase | ko00500//Starch and sucrose metabolism |
| PA2166 | PA2166 | 2.361 | Protein of uncharacterised function (DUF3509) | - |
| PA2173a | PA2173a | 2.226 | Uncharacterised protein | - |
| PA2174 | PA2174 | 1.656 | Uncharacterised protein | - |
| PA2203 | PA2203 | -2.285 | amino acid permease | - |
| PA2204 | PA2204 | -2.799 | ABC transporter | - |
| PA2213 | PA2213 | 1.463 | porin | - |
| PA2217 | PA2217 | -1.644 | aldehyde dehydrogenase | ko01220//Degradation of aromatic compounds;ko00930//Caprolactam degradation |
| PA2225 | PA2225 | 1.565 | Uncharacterised protein | - |
| PA2259 | ptxS | -1.244 | transcriptional regulator PtxS | - |
| PA2274 | PA2274 | -1.440 | antibiotic biosynthesis monooxygenase family protein | - |
| PA2279 | arsC | -2.050 | low molecular weight phosphatase | - |
| PA2287 | PA2287 | -1.249 | Uncharacterised protein | - |
| PA2312a | PA2312a | 2.133 | Uncharacterised protein | - |
| PA2314 | PA2314 | -2.174 | MFS transporter | - |
| PA2323 | PA2323 | -1.301 | glyceraldehyde-3-phosphate dehydrogenase | ko01200//Carbon metabolism;ko00010//Glycolysis / Gluconeogenesis;ko00030//Pentose phosphate pathway |
| PA2334 | PA2334 | 2.738 | transcriptional regulator | - |
| PA2350 | PA2350 | 1.458 | methionine import ATP-binding protein MetN 1 | ko02010//ABC transporters |
| PA2374 | PA2374 | 1.304 | MORN repeat variant family protein | - |
| PA2384 | PA2384 | -3.519 | ferric uptake regulator, Fur family | - |
| PA2386 | pvdA | -1.765 | L-ornithine N5-oxygenase | - |
| PA2391 | opmQ | -1.477 | probable outer membrane protein precursor | - |

| | | | | |
|---|---|---|---|---|
| PA2395 | pvdO | -1.553 | pyoverdine biosynthesis protein PvdO | - |
| PA2398 | fpvA | -1.342 | ferripyoverdine receptor | - |
| PA2404 | PA2404 | -1.225 | PA2404 | - |
| PA2407 | PA2407 | -1.676 | adhesion protein | ko02010//ABC transporters |
| PA2408 | PA2408 | -2.442 | ABC transporter ATP-binding protein | ko02010//ABC transporters |
| PA2409 | PA2409 | -3.708 | ABC transporter permease | ko02010//ABC transporters |
| PA2462 | PA2462 | -2.249 | hemagglutinin | - |
| PA2463 | PA2463 | -3.458 | hemolysin secretion/activation ShlB/FhaC/HecB family protein | - |
| PA2466 | foxA | -1.274 | ferrioxamine receptor FoxA | - |
| PA2470 | gtdA | -1.488 | gentisate 1,2-dioxygenase | ko00350//Tyrosine metabolism |
| PA2507 | catA | -1.461 | catechol 1,2-dioxygenase | ko00362//Benzoate degradation;ko01220//Degradation of aromatic compounds;ko00364//Fluorobenzoate degradation;ko00361//Chlorocyclohexane and chlorobenzene degradation;ko00623//Toluene degradation |
| PA2518 | xylX | 1.822 | toluate 1,2-dioxygenase subunit alpha | ko00362//Benzoate degradation;ko01220//Degradation of aromatic compounds;ko00364//Fluorobenzoate degradation;ko00622//Xylene degradation |
| PA2538 | PA2538 | 1.354 | Uncharacterised protein | - |
| PA2552 | PA2552 | -1.585 | acyl-CoA dehydrogenase | ko00281//Geraniol degradation |

| | | | | |
|---|---|---|---|---|
| PA2553 | PA2553 | -1.413 | acyl-CoA thiolase | ko02020//Two-component system;ko01200//Carbon metabolism;ko00620//Pyruvate metabolism;ko00630//Glyoxylate and dicarboxylate metabolism;ko01212//Fatty acid metabolism;ko00280//Valine, leucine and isoleucine degradation;ko00640//Propanoate metabolism;ko00650//Butanoate metabolism;ko00362//Benzoate degradation;ko00071//Fatty acid degradation;ko00380//Tryptophan metabolism;ko00310//Lysine degradation;ko00900//Terpenoid backbone biosynthesis;ko00072//Synthesis and degradation of ketone bodies |
| PA2557 | PA2557 | -1.407 | AMP-binding protein | - |
| PA2567 | PA2567 | -1.534 | sensor domain-containing phosphodiesterase | - |
| PA2607 | PA2607 | 1.226 | DsrH family protein | ko04122//Sulfur relay system |
| PA2652 | PA2652 | -2.108 | methyl-accepting chemotaxis protein | ko02020//Two-component system;ko02030//Bacterial chemotaxis |
| PA2675 | PA2675 | 1.909 | type II secretion system protein | ko03070//Bacterial secretion system |
| PA2701 | PA2701 | 1.302 | major facilitator superfamily transporter | - |
| PA2714 | PA2714 | -1.419 | molybdopterin oxidoreductase | - |
| PA2746a | PA2746a | 1.600 | predicted protein | - |
| PA2752 | PA2752 | 2.230 | Inner membrane protein YqaA | - |
| PA2754a | PA2754a | -2.690 | filamentous hemagglutinin | - |
| PA2766 | PA2766 | 1.394 | transcriptional regulator | - |
| PA2773 | PA2773 | 1.970 | membrane protein | - |

| | | | | |
|---|---|---|---|---|
| PA2788 | PA2788 | -2.143 | chemotaxis transducer | - |
| PA2862 | lipA | -1.686 | lactonizing lipase | ko00561//Glycerolipid metabolism |
| PA2867 | PA2867 | -3.315 | chemotaxis transducer | - |
| PA2879 | PA2879 | -1.544 | transcriptional regulator | - |
| PA2887 | atuB | -2.299 | citronellol catabolism dehydrogenase | ko00281//Geraniol degradation |
| PA2890 | atuE | -4.298 | isohexenylglutaconyl-CoA hydratase | ko00281//Geraniol degradation |
| PA2928 | PA2928 | 1.633 | Uncharacterised protein | - |
| PA3009 | PA3009 | 1.390 | ABC transporter ATP-binding protein | - |
| PA3038 | PA3038 | -2.391 | porin | - |
| PA3039 | PA3039 | 1.993 | transporter | - |
| PA3067 | PA3067 | 1.814 | transcriptional regulator | - |
| PA3079 | PA3079 | -1.889 | patched family protein | - |
| PA3126 | ibpA | -1.849 | heat-shock protein IbpA | - |
| PA3136 | PA3136 | 1.398 | secretion protein | - |
| PA3181 | PA3181 | -1.330 | keto-hydroxyglutarate-aldolase/keto-deoxy-phosphogluconate aldolase | ko01200//Carbon metabolism;ko00630//Glyoxylate and dicarboxylate metabolism;ko00030//Pentose phosphate pathway |
| PA3182 | pgl | -2.220 | 6-phosphogluconolactonase | ko01200//Carbon metabolism;ko00030//Pentose phosphate pathway |
| PA3183 | zwf | -2.022 | glucose-6-phosphate 1-dehydrogenase | ko01200//Carbon metabolism;ko00030//Pentose phosphate pathway;ko00480//Glutathione metabolism |
| PA3186 | oprB | -3.460 | porin B | - |
| PA3187 | PA3187 | -2.904 | ABC transporter ATP-binding protein | ko02010//ABC transporters |
| PA3189 | PA3189 | -1.454 | sugar ABC transporter permease | ko02010//ABC transporters |
| PA3190 | PA3190 | -1.862 | sugar ABC transporter substrate-binding protein | ko02010//ABC transporters |
| PA3192 | gltR | -1.969 | two-component response regulator GltR | - |

| | | | | |
|---|---|---|---|---|
| PA3194 | edd | -1.813 | phosphogluconate dehydratase | ko01200//Carbon metabolism;ko00030//Pentose phosphate pathway |
| PA3195 | gapA | -1.919 | glyceraldehyde 3-phosphate dehydrogenase | ko01230//Biosynthesis of amino acids;ko01200//Carbon metabolism;ko00010//Glycolysis / Gluconeogenesis |
| PA3209 | PA3209 | 12.001 | zinc/iron-chelating domain-containing protein | - |
| PA3232 | PA3232 | -1.374 | DNA polymerase III subunit epsilon | ko00230//Purine metabolism;ko00240//Pyrimidine metabolism;ko03440//Homologous recombination;ko03430//Mismatch repair;ko03030//DNA replication |
| PA3233 | PA3233 | -1.808 | cyclic nucleotide-binding domain protein | - |
| PA3234 | PA3234 | -3.020 | acetate permease | - |
| PA3235 | PA3235 | -2.130 | membrane protein | - |
| PA3236 | PA3236 | -1.907 | glycine betaine-binding protein | ko02010//ABC transporters |
| PA3282 | PA3282 | -2.139 | | - |
| PA3291 | PA3291 | -1.644 | DUF3304 domain-containing protein | - |
| PA3292 | PA3292 | -1.312 | DUF3304 domain-containing protein | - |
| PA3329 | PA3329 | -1.329 | Malonyl CoA-acyl carrier protein transacylase | - |
| PA3330 | PA3330 | -1.262 | short-chain dehydrogenase | - |
| PA3353 | PA3353 | -1.538 | pilus assembly protein PilZ | - |
| PA3358 | PA3358 | -11.948 | EamA family transporter | - |
| PA3376 | PA3376 | -2.276 | phosphonate C-P lyase system protein PhnK | - |
| PA3385 | amrZ | -1.372 | alginate and motility regulator Z | - |
| PA3425 | PA3425 | 1.219 | cupin | - |
| PA3442 | PA3442 | -1.243 | aliphatic sulfonates ABC transporter ATP-binding subunit | ko02010//ABC transporters;ko00920//Sulfur metabolism |

| | | | | |
|---|---|---|---|---|
| PA3444 | PA3444 | -1.421 | alkanesulfonate monooxygenase | ko00920//Sulfur metabolism |
| PA3484 | PA3484(Tse3) | -1.459 | Uncharacterized protein PAE221_01533 | - |
| PA3487 | pldA | -1.255 | phospholipase D | - |
| PA3502 | PA3502 | -2.117 | Uncharacterised protein | - |
| PA3506 | PA3506 | -1.850 | probable decarboxylase | ko01230//Biosynthesis of amino acids;ko00650//Butanoate metabolism;ko01210//2-Oxocarboxylic acid metabolism;ko00770//Pantothenate and CoA biosynthesis;ko00290//Valine, leucine and isoleucine biosynthesis;ko00660//C5-Branched dibasic acid metabolism |
| PA3507 | PA3507 | -2.411 | short-chain dehydrogenase | - |
| PA3508 | PA3508 | -2.018 | transcriptional regulator | - |
| PA3510 | PA3510 | -1.481 | cupin domain protein | - |
| PA3519 | PA3519 | -1.509 | Pyrroloquinoline quinone (Coenzyme PQQ) biosynthesis protein C | - |
| PA3523 | PA3523 | -1.438 | resistance-nodulation-cell division (RND) efflux membrane fusion protein | - |
| PA3526 | PA3526 | -2.106 | probable outer membrane protein precursor | - |
| PA3568 | PA3568 | -3.847 | propionyl-CoA synthetase | ko00640//Propanoate metabolism |
| PA3569 | mmsB | -2.780 | 3-hydroxyisobutyrate dehydrogenase | ko00280//Valine, leucine and isoleucine degradation |
| PA3570 | mmsA | -2.312 | methylmalonate-semialdehyde dehydrogenase | ko01200//Carbon metabolism;ko00280//Valine, leucine and isoleucine degradation;ko00640//Propanoate metabolism;ko00410//beta-Alanine metabolism;ko00562//Inositol phosphate |

| | | | | |
|---|---|---|---|---|
| PA3593 | PA3593 | 1.591 | acyl-CoA dehydrogenase | - |
| PA3601 | PA3601 | -2.114 | 50S ribosomal protein L31 | ko03010//Ribosome |
| PA3612 | PA3612 | 1.463 | Putative cytoplasmic protein | - |
| PA3662 | PA3662 | -3.114 | Uncharacterised protein | - |
| PA3714 | PA3714 | -1.513 | LuxR family transcriptional regulator | ko02020//Two-component system |
| PA3720 | PA3720 | -1.775 | Uncharacterised protein | - |
| PA3721 | nalC | -1.702 | transcriptional regulator | - |
| PA3722 | PA3722 | -2.087 | Uncharacterised protein | - |
| PA3727 | PA3727 | -1.526 | nuclease | - |
| PA3728 | PA3728 | -1.759 | AAA domain family protein | - |
| PA3729 | PA3729 | -1.637 | Inner membrane protein YqiK | - |
| PA3731 | PA3731 | -1.209 | phage shock protein A | - |
| PA3740 | PA3740 | -1.216 | Type II secretory pathway, ATPase PulE/Tfp pilus assembly pathway, ATPase PilB | - |
| PA3745 | rpsP | 1.231 | 30S ribosomal protein S16 | ko03010//Ribosome |
| PA3758 | PA3758 | -1.434 | N-acetylglucosamine-6-phosphate deacetylase | ko00520//Amino sugar and nucleotide sugar metabolism |
| PA3759 | PA3759 | -1.294 | aminotransferase | ko00250//Alanine, aspartate and glutamate metabolism;ko00520//Amino sugar and nucleotide sugar metabolism |
| PA3760 | PA3760 | -1.408 | phosphoenolpyruvate--protein phosphotransferase | - |
| PA3761 | PA3761 | -1.296 | N-acetyl-D-glucosamine phosphotransferase system transporter | ko00520//Amino sugar and nucleotide sugar metabolism;ko02060//Phosphotransferase system (PTS) |
| PA3838 | PA3838 | -1.475 | ABC transporter ATP-binding protein | - |
| PA3842 | PA3842 | 1.216 | chaperone | - |
| PA3856 | PA3856 | 1.254 | ketosteroid isomerase | - |
| PA3867 | PA3867 | 1.511 | DNA invertase | - |

| | | | | |
|---|---|---|---|---|
| PA3870 | moaA1 | 1.329 | molybdenum cofactor biosynthesis protein A | ko00790//Folate biosynthesis;ko04122//Sulfur relay system |
| PA3871 | PA3871 | 1.444 | peptidyl-prolyl cis-trans isomerase | - |
| PA3873 | narJ | 1.417 | respiratory nitrate reductase subunit delta | ko02020//Two-component system;ko00910//Nitrogen metabolism |
| PA3874 | narH | 1.383 | respiratory nitrate reductase subunit beta | ko02020//Two-component system;ko00910//Nitrogen metabolism |
| PA3875 | narG | 1.476 | respiratory nitrate reductase subunit alpha | ko02020//Two-component system;ko00910//Nitrogen metabolism |
| PA3876 | narK2 | 1.645 | nitrite extrusion protein 2 | ko00910//Nitrogen metabolism |
| PA3884 | PA3884 | 12.097 | putative membrane protein | - |
| PA3923 | PA3923 | -1.339 | adhesin | - |
| PA3935 | tauD | -1.871 | taurine dioxygenase | ko00920//Sulfur metabolism;ko00430//Taurine and hypotaurine metabolism |
| PA3937 | PA3937 | -1.929 | taurine ABC transporter ATP-binding protein | ko02010//ABC transporters;ko00920//Sulfur metabolism |
| PA4022 | PA4022 | -1.474 | aldehyde dehydrogenase | ko00620//Pyruvate metabolism;ko00010//Glycolysis / Gluconeogenesis |
| PA4023 | PA4023 | -2.337 | transporter | - |
| PA4038 | PA4038 | 1.567 | possible ABC transporter permease component | - |
| PA4072 | PA4072 | -1.542 | amino acid permease | - |
| PA4073 | PA4073 | -1.489 | Phenylacetaldehyde dehydrogenase | ko00360//Phenylalanine metabolism;ko00643//Styrene degradation |
| PA4089 | PA4089 | 2.707 | 3-ketoacyl-ACP reductase | ko01212//Fatty acid metabolism;ko00061//Fatty acid biosynthesis;ko00780//Biotin metabolism;ko01040//Biosynthesis of unsaturated fatty acids |

| | | | | |
|---|---|---|---|---|
| PA4093 | PA4093 | -2.515 | thioesterase | - |
| PA4107 | PA4107 | -2.520 | EF-hand domain pair family protein | - |
| PA4124 | hpcB | -1.495 | 3,4-dihydroxyphenylacetate 2,3-dioxygenase | ko01220//Degradation of aromatic compounds;ko00350//Tyrosine metabolism |
| PA4139 | PA4139 | -1.531 | Uncharacterized protein PAE221_02180 | - |
| PA4140 | PA4140 | -1.620 | FAD-linked oxidase | - |
| PA4151 | acoB | 1.356 | acetoin catabolism protein AcoB | ko01200//Carbon metabolism;ko00620//Pyruvate metabolism;ko00010//Glycolysis / Gluconeogenesis;ko00020//Citrate cycle (TCA cycle) |
| PA4171 | PA4171 | 1.501 | glutamine amidotransferase | - |
| PA4188 | PA4188 | -1.842 | dihydrodipicolinate synthase family protein | ko01230//Biosynthesis of amino acids;ko00300//Lysine biosynthesis;ko00261//Monobactam biosynthesis |
| PA4189 | PA4189 | -2.000 | aldehyde dehydrogenase | ko00330//Arginine and proline metabolism |
| PA4191 | PA4191 | -1.353 | iron oxidase | - |
| PA4192 | PA4192 | -2.193 | ABC transporter ATP-binding protein | - |
| PA4198 | PA4198 | -1.273 | acyl-CoA synthetase | - |
| PA4204 | ppgL | -1.214 | gluconolactonase PpgL | - |
| PA4211 | phzB1 | -1.678 | phenazine biosynthesis protein phzB 1 | - |
| PA4213 | phzD1 | -1.698 | phenazine biosynthesis protein PhzD | - |
| PA4214 | phzE1 | -1.324 | phenazine biosynthesis protein PhzE | - |
| PA4215 | phzF1 | -1.369 | 2,3-dihydro-3-hydroxyanthranilate isomerase | - |
| PA4221 | fptA | -1.568 | Fe(III)-pyochelin outer membrane receptor | - |
| PA4222 | PA4222 | -1.774 | ABC transporter ATP-binding protein | - |
| PA4223 | PA4223 | -2.059 | ABC transporter ATP-binding protein | ko02010//ABC transporters |
| PA4224 | pchG | -1.551 | pyochelin biosynthetic protein PchG | ko01053//Biosynthesis of siderophore group nonribosomal peptides |

| | | | | |
|---|---|---|---|---|
| PA4225 | pchF | -2.527 | pyochelin synthetase | ko01053//Biosynthesis of siderophore group nonribosomal peptides |
| PA4226 | pchE | -2.000 | dihydroaeruginoic acid synthetase | ko01053//Biosynthesis of siderophore group nonribosomal peptides |
| PA4231 | pchA | -1.843 | salicylate biosynthesis isochorismate synthase | ko00130//Ubiquinone and other terpenoid-quinone biosynthesis;ko01053//Biosynthesis of siderophore group nonribosomal peptides |
| PA4290 | PA4290 | -1.329 | chemotaxis transducer | ko02020//Two-component system;ko02030//Bacterial chemotaxis |
| PA4291 | PA4291 | 1.270 | hypothetical protein | - |
| PA4298 | PA4298 | -1.629 | Protein of uncharacterised function (DUF3613) | - |
| PA4306 | flp | -1.410 | type IVb pilin Flp | - |
| PA4309 | pctA | -1.419 | methyl-accepting chemotaxis protein PctA | ko02020//Two-component system;ko02030//Bacterial chemotaxis |
| PA4310 | pctB | -1.687 | methyl-accepting chemotaxis protein PctB | ko02020//Two-component system;ko02030//Bacterial chemotaxis |
| PA4324 | PA4324 | -2.674 | pilus assembly protein PilZ | - |
| PA4326 | PA4326 | -2.200 | lipoprotein | - |
| PA4332 | PA4332 | -1.419 | FOG: GGDEF domain | - |
| PA4500 | PA4500 | -1.586 | ABC transporter | ko02010//ABC transporters;ko02030//Bacterial chemotaxis |
| PA4501 | opdP | -2.560 | porin | - |
| PA4502 | PA4502 | -2.093 | ABC transporter | ko02010//ABC transporters;ko02030//Bacterial chemotaxis |
| PA4504 | PA4504 | -1.624 | ABC transporter permease | ko02010//ABC transporters |
| PA4505 | PA4505 | -1.528 | ABC transporter ATP-binding protein | ko02010//ABC transporters |
| PA4506 | PA4506 | -1.516 | peptide ABC transporter ATP-binding protein | ko02010//ABC transporters |

| | | | | |
|---|---|---|---|---|
| PA4520 | PA4520 | -1.391 | chemotaxis transducer | ko02020//Two-component system;ko02030//Bacterial chemotaxis |
| PA4563 | rpsT | 1.423 | 30S ribosomal protein S20 | ko03010//Ribosome |
| PA4571 | PA4571 | -2.853 | cytochrome c | - |
| PA4573 | PA4573 | 1.360 | cation transporter | - |
| PA4582 | PA4582 | -1.434 | SPFH domain / Band 7 family protein | - |
| PA4590 | pra | -1.482 | protein activator | - |
| PA4592 | PA4592 | -1.258 | probable outer membrane protein precursor | - |
| PA4596 | PA4596 | -3.429 | transcriptional regulator | - |
| PA4625 | PA4625 | -1.401 | hemagglutination protein | - |
| PA4633 | PA4633 | -2.364 | chemotaxis protein | ko02020//Two-component system;ko02030//Bacterial chemotaxis |
| PA4644 | PA4644 | 1.251 | Tryptophan synthase beta chain like | - |
| PA4675 | PA4675 | -1.497 | TonB-dependent receptor | - |
| PA4683 | PA4683 | -2.179 | Uncharacterized protein PAE221_00037 | - |
| PA4709 | PA4709 | 1.355 | hemin degrading factor | - |
| PA4746 | PA4746 | 1.260 | ribosome maturation factor RimP | - |
| PA4764 | fur | 1.299 | ferric uptake regulation protein | - |
| PA4766 | PA4766 | 2.354 | protein RnfH | - |
| PA4768 | smpB | 1.411 | SsrA-binding protein | - |
| PA4780 | PA4780 | 1.611 | serine protein kinase RIO | - |
| PA4789 | PA4789 | 1.534 | pyrophosphatase | - |
| PA4817 | PA4817 | 2.426 | zinc/iron-chelating domain-containing protein | - |
| PA4843 | PA4843 | -4.397 | diguanylate cyclase response regulator | - |
| PA4844 | PA4844 | -2.290 | chemotaxis transducer | - |
| PA4889 | PA4889 | 1.261 | oxidoreductase | - |
| PA4901 | mdlC | -1.512 | benzoylformate decarboxylase | ko00627//Aminobenzoate degradation |
| PA4911 | PA4911 | -1.297 | branched-chain amino acid ABC transporter permease | ko02010//ABC transporters |

| | | | | |
|---|---|---|---|---|
| PA4919 | pncB1 | -1.621 | nicotinate phosphoribosyltransferase | ko00760//Nicotinate and nicotinamide metabolism |
| PA4929 | PA4929 | -1.840 | sensor domain-containing diguanylate cyclase | - |
| PA4989 | PA4989 | -1.234 | transcriptional regulator | - |
| PA5054 | hslU | -1.298 | ATP-dependent protease ATP-binding subunit HslU | - |
| PA5072 | PA5072 | -1.354 | chemotaxis transducer | ko02020//Two-component system;ko02030//Bacterial chemotaxis |
| PA5098 | hutH | -1.932 | histidine ammonia-lyase | ko00340//Histidine metabolism |
| PA5099 | PA5099 | -1.373 | transporter | - |
| PA5100 | hutU | -1.668 | urocanate hydratase | ko00340//Histidine metabolism |
| PA5153 | PA5153 | -1.940 | amino acid ABC transporter substrate-binding protein | - |
| PA5154 | PA5154 | -1.853 | ABC transporter permease | - |
| PA5155 | PA5155 | -1.606 | amino acid ABC transporter permease | - |
| PA5167 | PA5167 | -1.207 | C4-dicarboxylate-binding protein | ko02020//Two-component system |
| PA5181 | PA5181 | -1.343 | oxidoreductase | - |
| PA5285 | PA5285 | 1.320 | Uncharacterised protein | - |
| PA5289 | PA5289 | 1.518 | membrane fusogenic activity family protein | - |
| PA5383 | PA5383 | -1.707 | membrane protein | - |
| PA5387 | cdhC | 2.967 | carnitine dehydrogenase | - |
| PA5396 | PA5396 | -1.535 | membrane dipeptidase family protein | - |
| PA5397 | PA5397 | -2.565 | hydrocarbon binding protein | - |
| PA5398 | dgcA | -2.146 | dimethylglycine catabolism protein DgcA | - |
| PA5399 | dgcB | -1.630 | (Fe-S)-binding protein | - |
| PA5410 | gbcA | -1.560 | Rieske (2Fe-2S) protein | - |

| | | | | |
|---|---|---|---|---|
| PA5415 | glyA1 | -2.044 | serine hydroxymethyltransferase | ko01230//Biosynthesis of amino acids;ko01200//Carbon metabolism;ko00630//Glyoxylate and dicarboxylate metabolism;ko00260//Glycine, serine and threonine metabolism;ko00680//Methane metabolism;ko00670//One carbon pool by folate;ko00460//Cyanoamino acid metabolism |
| PA5416 | soxB | -2.042 | sarcosine oxidase subunit beta | ko00260//Glycine, serine and threonine metabolism |
| PA5417 | soxD | -2.744 | sarcosine oxidase subunit delta | ko00260//Glycine, serine and threonine metabolism |
| PA5418 | soxA | -2.625 | sarcosine oxidase subunit alpha | ko00260//Glycine, serine and threonine metabolism |
| PA5419 | soxG | -1.468 | sarcosine oxidase subunit gamma | ko00260//Glycine, serine and threonine metabolism |
| PA5420 | purU2 | -1.969 | formyltetrahydrofolate deformylase | ko00630//Glyoxylate and dicarboxylate metabolism;ko00670//One carbon pool by folate |
| PA5421 | fdhA | -1.876 | glutathione-independent formaldehyde dehydrogenase | ko01200//Carbon metabolism;ko00680//Methane metabolism;ko00625//Chloroalkane and chloroalkene degradation |
| PA5444 | PA5444 | 1.566 | transporter | - |
| PA5445 | PA5445 | -1.549 | coenzyme A transferase | ko01200//Carbon metabolism;ko00620//Pyruvate metabolism;ko00650//Butanoate metabolism;ko00020//Citrate cycle (TCA cycle) |
| PA5462 | PA5462 | 1.895 | Holliday junction resolvase | - |
| PA5491 | PA5491 | 1.240 | cytochrome | - |
| PA5515 | PA5515 | 1.324 | Protein of uncharacterised function (DUF3301) | - |

| | | | | |
|---|---|---|---|---|
| PA5536 | PA5536 | 2.970 | DksA/TraR family C4-type zinc finger protein | - |
| PA5550 | glmR | 1.377 | GlmR transcriptional regulator | - |
| PA5566 | PA5566 | -1.603 | hypothetical protein | - |

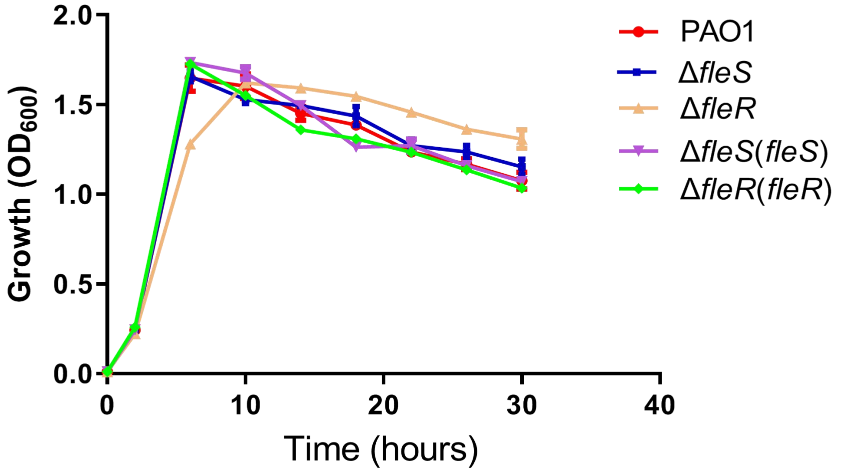


**Fig. S1.** Growth of PAO1 and its *fleS* and *fleR* mutants as well as corresponding complemented strains. The data is the mean of five replicates with standard deviations.

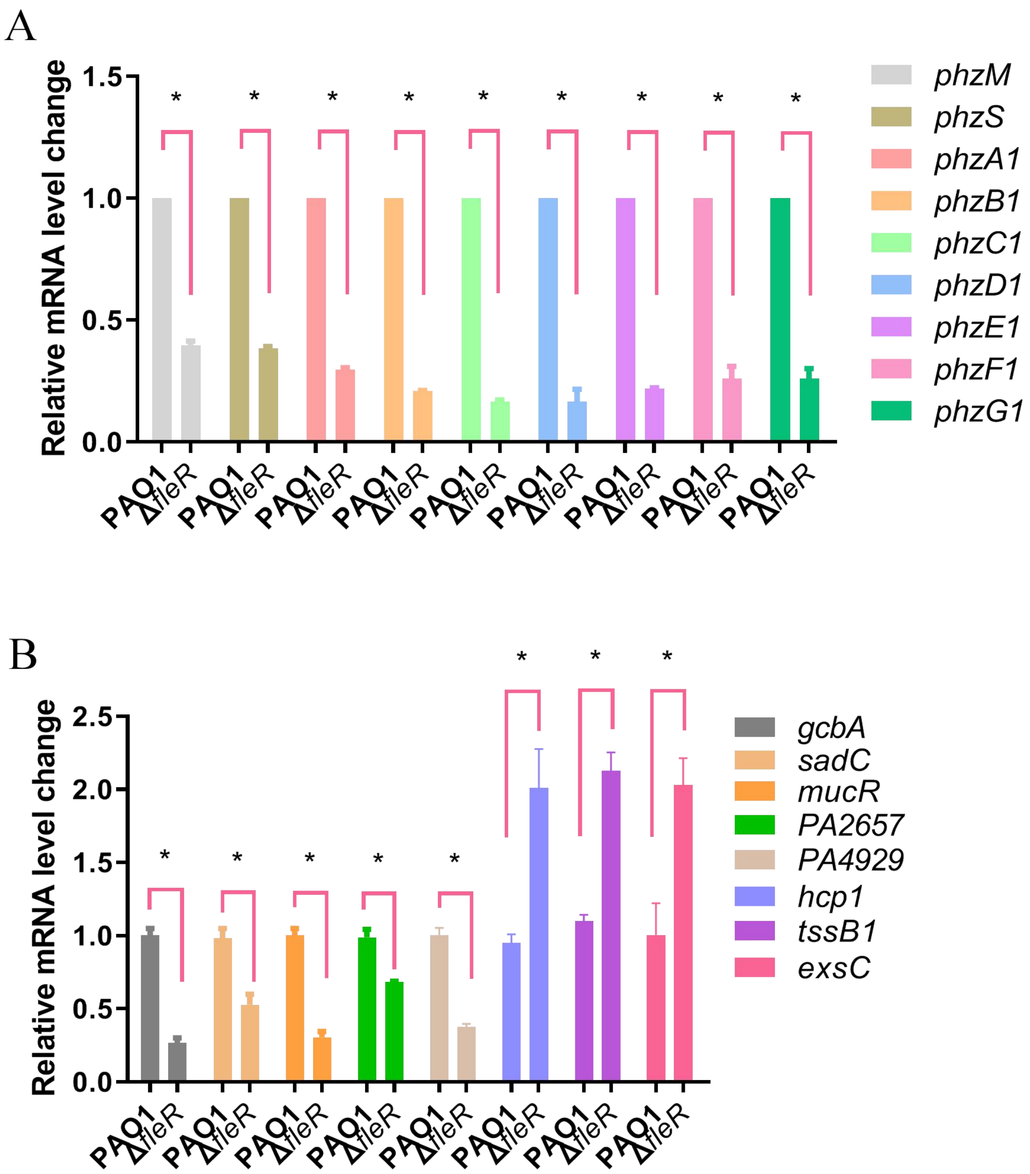


**Fig. S2.** The relative mRNA levels of the genes involved in pyocyanin biosynthesis (*phzMS*, *phzA1-G1*) (A), c-di-GMP metabolism (*gcbA*, *sadC*, *mucR*, *PA2657*, *PA4929*) and secretion systems (*hcp1*, *tssB1*, *exsC*) (B) in PAO1 and Δ*fleR*. The data is the mean of triplicates with standard deviations. *: $p < 0.05$, ns: not significant, tested by Student's *t*-test.

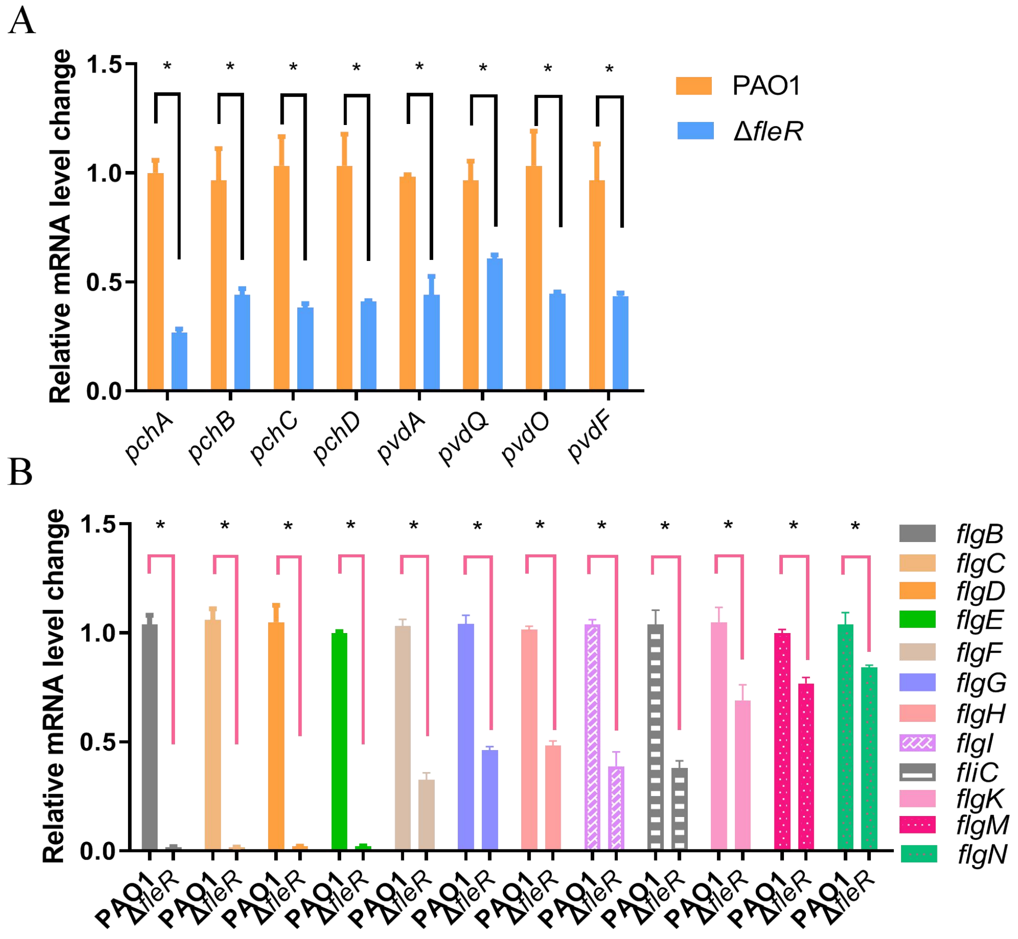


**Fig. S3.** The relative mRNA levels of the genes encoding pyoverdine (A) (*pvdA, pvdQ, pvdO, pvdF*), pyochelin (*pchABCD*) and flagellum biosynthesis (B) (*flgB-I*, *fliC*, *flgK*, *flgMN*) in PAO1 and Δ*fleR*. The data is the mean of triplicates with standard deviations. *: $p < 0.05$, ns: not significant, tested by Student's *t*-test.

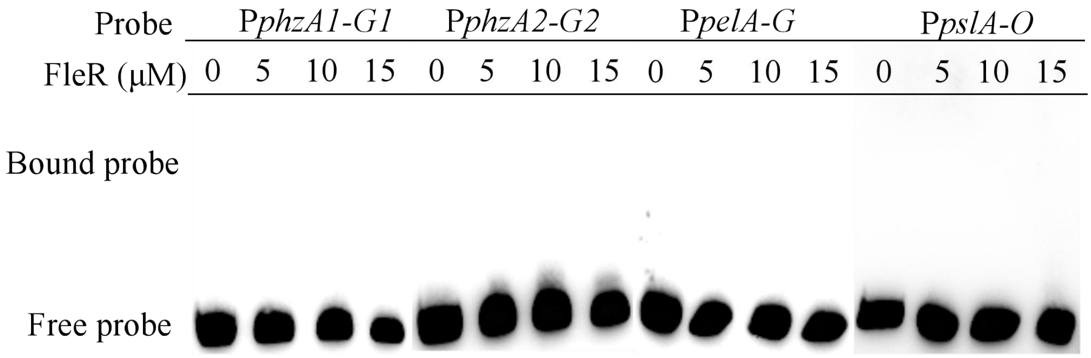


**Fig. S4.** EMSA examinations of FleR binding to the promoters of pyocyanin biosynthesis operons *phzA1-G1* (P*phzA1-G1*) and *phzA2-G2* (P*phzA2-G2*), Pel biosynthesis operon *pelA-G* (P*pelA-G*) and Psl biosynthesis operon *pslA-O* (*pslA-O*).

```
flgBCDE     GATCATTCGCCACGAGTGAGCCGTGTCCCCGGCGGCCGGAAGCATGGGAAAGGCC  55
flgFGHIJKL  ................CAGACCGAGGATGCGGTGACCCAGACCATCATCAACCTG  39
fliC        ...................ACGGGAGGGCTAAAGAAAATCGCCGGGGGGTCGATG  36
fleSR       ................AAGGCCTGGACCTCAAGGACTA....CCTGGCCAACCTC  35
Consensus                       c            g        c

flgBCDE     CGATAT.CGAC.GATATCGGGCCTTTTTTCGTTT..GCCGGAGAAG..CCGTTCC  104
flgFGHIJKL  CGCTGA.TGAC.GGCGTGCCGGTGCCGTCCGGGC..ACCGGCAAGG..CCCTTGC  88
fliC        CAATGGGTGTCGGAACTTCCACCCTCTGCCGGACCAACCGGGGGCGGTTCAGGAC  91
fleSR       GAGCAG.GGCCTGATCCAGCAGGCCCTCGACGAC..GCCGGCGGAG..TGGTCGC  85
Consensus           g c g                        ccgg    g        c

flgBCDE     AGGGGCTTGCCACCCTTGCC...GGCAAGCGGAAGCGGCTTGCCGCTTGCCGCTT  156
flgFGHIJKL  CGCCAGCGGCAACCCCTGGCCAGGGGTTGCCCCCGAAGCCTCCGAAAGGAGGCTT  143
fliC        CGATATTGGCGAGTCCTCTTCGAAGCATGTAACCCACTGAAGAGGAAGAGAAAAA  146
fleSR       GCGGGCCGCCGAACGCCTGC..GCATCCGCCGCACCACGCTGGTAGAGAAGATGC  138
Consensus            c a                g

flgBCDE     CCGGGTTACCCTTAAATAAACTCAAGTTATTGAATATAAAGGTTTTTAATAATTT  211
flgFGHIJKL  TCTGTTT..TATAAAATCCTTTTAAATCAATGTATTAATTGTTTTTTCGAATTCT  196
fliC        GAAAATGTTGATT..TTTTCTCTAAAGCTCCGCCGGGAAACGCCGATAAACACCA  199
fleSR       GCAAGTACGGCATGAGCCGGCGTGA..CGACGACCTGTCGGATGATTGACAGGTC  191
Consensus        t                  a      g              t

flgBCDE     GGCACGGGCCTTGCTGAAGTGTTGGCG.ACGAAAGCCACACCGGCAAACAGGTTC  265
flgFGHIJKL  GGCACGGCGCTTGCTGGATAACCTGCA.A.GAAAGCC....CGGCGGATTGTGCC  245
fliC        TGAACGCGAATTCTTGGGGCACCTGAGCAAGCAGGCCGAGAGATCGCAAGCTCAG  254
fleSR       GTTTCGCAACGCTTTGATTTTCAAATG.AAAAAAATTTAGGCA.CGGGTATTGCT  244
Consensus       cg        tg            a   a           c

flgBCDE     GCAGCCATGA.GCATCAGTTTCGACAGA.GCACTCGGTATCCACCAGCAG.....  313
flgFGHIJKL  GGTGCCTGGA.GGATTCGATGGACAAGATGCTGTACGTCTCCATGAGCGGGGCCA  299
fliC        GTAACCGAAATAGGTCCTTTGGAGGAAATCACCATGGCCCTTACAGTCAACACGA  309
fleSR       ATATCTCCGT.CGACCGACAGAACCATGACGTCGCGCCGACCGAGAGAAACGCAA  298
Consensus       c                    a

flgBCDE     .................................                        313
flgFGHIJKL  GCCAGAACACCCTGGCGATGCGTGCTCATGCCA                        332
fliC        ACA..............................                        312
fleSR       TGCAACCAGC.......................                        308
Consensus
```

**Fig. S5.** The promoters of *flgBCDE*, *flgFGHIJKL*, *fliC* and *fleSR* are aligned using clustal X. Blue color represents 100 % identity and pink color represents 75 % identity.
